# SpinX: Time-resolved 3D Analysis of Mitotic Spindle Dynamics using Deep Learning Techniques and Mathematical Modelling

**DOI:** 10.1101/2021.09.21.461203

**Authors:** David Dang, Christoforos Efstathiou, Dijue Sun, Nishanth Sastry, Viji M. Draviam

## Abstract

Time-lapse microscopy movies have transformed the study of subcellular dynamics. However, manual analysis of movies can introduce bias and variability, obscuring important insights. While automation can overcome such limitations, spatial and temporal discontinuities in time-lapse movies render methods such as object segmentation and tracking difficult. Here we present SpinX, a framework for reconstructing gaps between successive frames by combining Deep Learning and mathematical object modelling. By incorporating expert feedback through selective annotations, SpinX identifies subcellular structures, despite confounding neighbour-cell information, non-uniform illumination and variable marker intensities. The automation and continuity introduced allows precise 3-Dimensional tracking and analysis of spindle movements with respect to the cell cortex for the first time. We demonstrate the utility of SpinX using distinct spindle markers and drug treatments. In summary, SpinX provides an exciting opportunity to study spindle dynamics in a sophisticated way, creating a framework for step changes in studies using time-lapse microscopy.

## Introduction

Computational image analysis tools and single-cell imaging methods can accelerate cell biology studies^1–3^ and drug discovery efforts^4^. Although Deep Learning (DL) has revolutionised the automated and unbiased analysis of still microscopy images for object identification^5–8^, it has not been successfully extended to time-lapse movies for analysing sub-cellular structural dynamics through time and 3-Dimensional (3D) space. Extending DL to time-lapse movies has faced at least two critical hurdles: first, the precise tracking of structures through time requires tailored 3D object modelling tools to overcome spatial and temporal discontinuities that are intrinsic to time-lapse movies. Second, feature-rich analysis supported by DL methods require large volumes of high-resolution time-lapse movie datasets^9^. Nevertheless, as DL architectures for still images of fixed-cells^10–12^ have helped overcome the drawback of manual analysis (with respect to image segmentation which is inherently tedious, slow and error prone), developing new DL architectures for time-lapse movies of live-cells can advance quantitative 3D studies of subcellular, cellular and tissue level dynamics.

Automated tools to analyse dynamic changes in intensities captured in live-cell movies are available^2,13^, but those that can reliably track precise changes in 3D shape and motion are challenging. Spatial and temporal discontinuities in time-lapse movies prevent the continuous tracking of objects in motion, and these are most common in cases where phototoxicity or photobleaching are predominant factors^14^. For example, fluorescent labelling of dividing cells with condensed chromosomes is well known to induce phototoxicity^15^, rendering it difficult to continuously image and track the dynamic movements of subcellular structures during mitosis.

The mitotic spindle is a complex and dynamic structure that is dependent on the function and regulation of multiple factors: the microtubule cytoskeleton^16^, molecular motors^17–19^, actin clouds^20^, cell cortex rigidity^21^, cell-cell adhesion complexes^22,23^ and chromosome congression^24^. The spindle undergoes complex 3D movements in longitudinal, equatorial and axial directions, by integrating both intracellular and extracellular cues^25–29^ that ultimately guide the spindle’s position to precisely define the future plane of cell division^30,31^. Thus, the complex movements of the mitotic spindle makes it an ideal subcellular model for testing the efficacy of DL-based video analysis methods aimed at extracting reliable and dynamic 3D information. Conventional computational methods^32–35^ have not been successful in continuous automated tracking of spindle largely due to the lack of spatio-temporal continuity in time-lapse movies.

As DL methods are data hungry^36^, we first generated a large dataset of high-resolution time-lapse movies of mitotic spindle movements in human epithelial cells expressing a fluorescent tagged microtubule marker protein, Tubulin. Using this large dataset of 28,350 images, we built a comprehensive and extensible computational framework, SpinX, which bridges the gaps between discontinuous frames in time-lapse movies by utilising state-of-the-art DL technologies and mathematical object modelling for 3D reconstruction of the mitotic spindle and cell cortex. Through stepwise benchmarking and detailed manual assessments, we demonstrate the potential of the 3D reconstruction module in overcoming spatiotemporal discontinuity in time-lapse movies of mitotic spindle and cell cortex. We establish the generalisation capacity of the SpinX framework for spindle segmentation using different microtubule-associated molecular markers and microscopy systems. Finally, using SpinX to track 3D movements of the spindle in cells treated with MARK2 kinase inhibitor, which has been used for *in vitro* but not *in vivo* studies, we showcase the application of AI-based time-lapse movie analysis in accelerating cytoskeletal research and drug development.

## Results

### Computational framework to track 3D movements of the mitotic spindle

To create a computational framework for accurately tracking the movements of the mitotic spindle in 3D, we first generated our own training dataset of high-resolution time-lapse movies for building the DL network. For this purpose, we labelled H2B-GFP (a chromosome marker) expressing cells^27^ with one of two different markers for the mitotic spindle, mCherry-Tubulin or SiR-Tubulin dye. Both markers have been established to decorate the microtubules of the mitotic spindle but with varying intensities^27,38^. The cell cortex was tracked label-free using brightfield images. A total of nearly five Terabytes of time-lapse movies were generated by imaging spindles in over hundreds of cells exposed to MG132 (a proteasome inhibitor to prevent metaphase-anaphase transition^39^). Three *z*-slice images (with a *z*-gap of 2 microns) were acquired once every three minutes to ensure that the spatiotemporal discontinuity in 3D time-lapse movies did not impair manual tracking of spindle movements. As expected, although live-cell movies are powerful in revealing dynamic cellular behaviour, they capture highly heterogeneous information across and within cells through time (Fig. 1a), making it difficult to quantitatively track spindle movements in 3D using traditional image segmentation methods^27^. We observed several challenges in segmenting live-cell imaging data using traditional image analysis tools: i) variability in sample illumination and protein expression between cells, where occasionally signal intensity can be highly uniform; ii) noise from neighbouring objects exacerbating low signal-to-noise ratios; and iii) loss of focus resulting in blurry images due to natural 3D movements over time (Fig. 1a). To overcome these challenges in tracking subcellular dynamics, we developed SpinX’s AI module by adapting the state-of-the-art Mask R-CNN DL architecture^37^ (see below and Fig. 1b).

**Figure 1.**
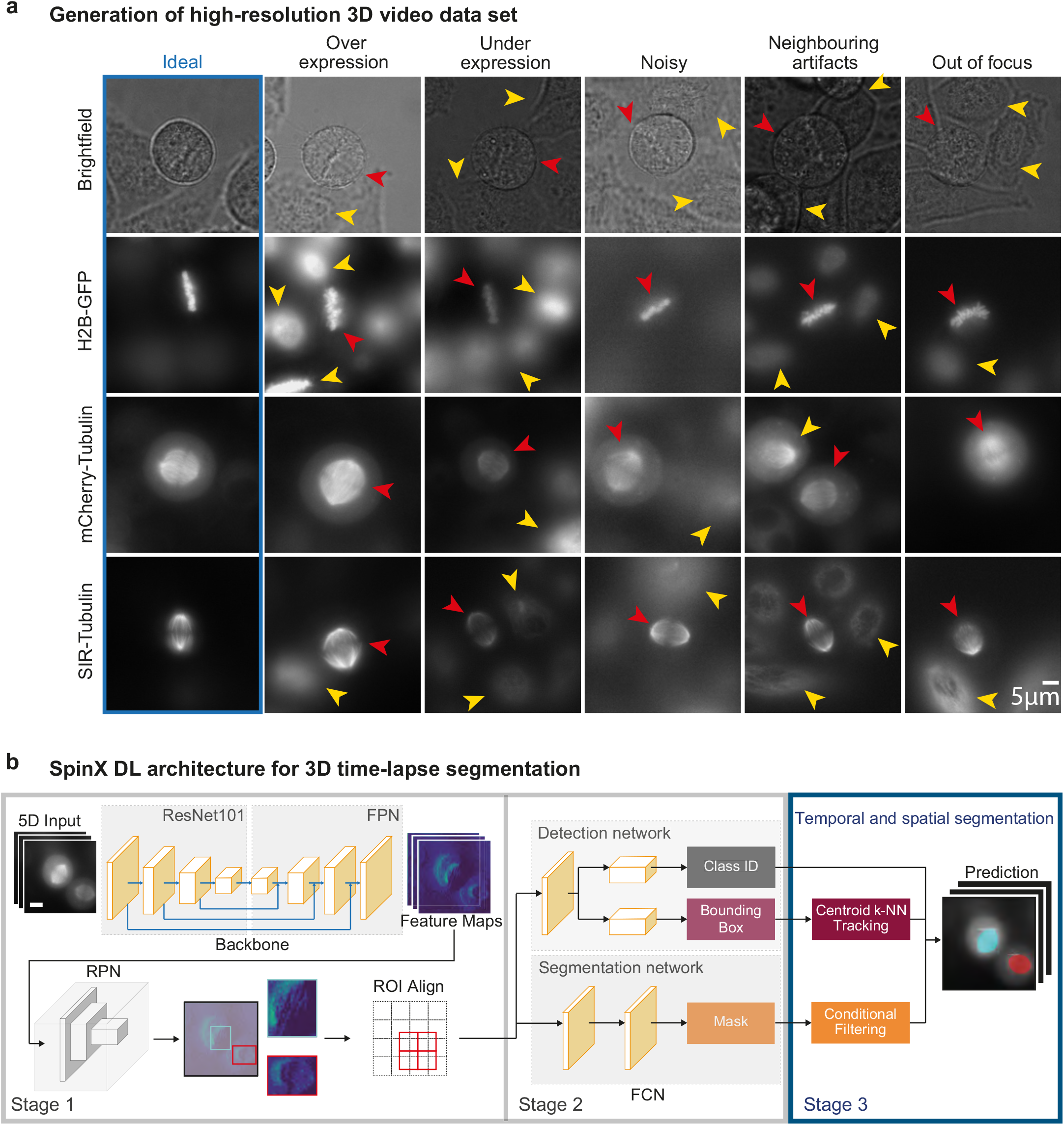
Video dataset and DL architecture for spindle and cell cortex image segmentation. **a**. Representative images show complex variation in illumination and marker intensities intrinsic to time-lapse movies of subcellular structures (cell cortex, brightfield (label-free); chromosomes, H2B-GFP; mitotic spindle, mCherry-Tubulin or SiR-Tubulin dye). Ideal images are shown within the blue box. Red and yellow arrowheads indicate the object of interest and interfering variation(s), respectively. Scale bar: 5 *µ*m. **b**. Deep Learning model architecture of SpinX. The model expands the pre-existing Mask R-CNN architecture (ResNet101, FPN, RPN, ROI Align, and FCN^37^), by introducing a third stage (blue box). In Stage 1, the network performs object detection followed by the segmentation of spindle and cell cortex in Stage 2. Stage 3 (highlighted in blue) links temporal and spatial information in 3D live-cell movies through tracking, and generates a consistent mask of the same object through time. The inputs of the model are grayscale or RGB images of various sizes (5D input). The outputs are binary masks of the same size as inputs with predicted foreground regions, bounding box coordinates (rectangular boxes in teal and red, Stage 1) and the corresponding Class ID (Stage 2). Scale bar: 10 *µ*m.

SpinX’s AI architecture identifies fluorescently-labelled spindles within label-free single-cell compartments by integrating three stages (Fig. 1b), where the first two stages closely resemble a ‘native’ Mask R-CNN-based DL architecture (for details see Methods). The first stage combines a convolutional backbone architecture - comprised of a Residual Network with 101 layers (ResNet101)^40^ and a Feature Pyramid Network (FPN)^41^ - with a Region Proposal Network (RPN)^42^. The aligned regions of interest (ROIs) are then passed onto the second stage of SpinX’s architecture: a Fully Convolutional Network (FCN)^43^ that simultaneously performs object classification and segmentation for every aligned ROI (Fig. 1b). The classes, bounding box information and masks generated through the first two stages of SpinX’s DL architecture are passed to the third and final stage consisting of two modules (see blue box Fig. 1b). In Stage 3, a ‘Conditional Filtering’ module filters and discards detected objects based on their i) confidence score that is derived from the prediction of the network; ii) associated pixel count (i.e. area) that eliminates any artefacts including objects much smaller than the spindle; and iii) location within the image canvas that allows the elimination of any detected object close to the image border as it would exhibit an incomplete shape. A second module in Stage 3, ‘Centroid *k*-NN Tracking’ exploits the bounding box information, wherein the predicted bounding box centroid coordinates are fed into a single point k-nearest neighbours algorithm (*k*-NN) for tracking a singular detected object through time.

SpinX’s workflow is comprehensive, including preprocessing of data with annotations to 3D modelling. For training, validation, and testing of the pipeline, we used a total of 2,180 label-free (brightfield) images to deduce the cell membrane (randomly selected from an image pool of 13,230 images) and 2,320 fluorescently- or dye-labelled microtubules images to deduce the mitotic spindle (randomly selected from an image pool of 15,120 images) (Supplementary Fig. 1). Annotation of our training dataset (n = 1300 images for cell cortex model; n = 1390 images for spindle model) was done automatically and subsequently corrected manually (See ‘Annotations’ in Methods). For automated label generation, we combined conventional image processing methods for specifically annotating chromosome and dye-based spindle (SiR-Tubulin) images (Supplementary Fig. 2). The rate of automated labelling was nearly 100-fold faster than manual labelling (Supplementary Fig. 3a and b). Our automated label generation pipeline correctly labelled chromosomes and dye-based spindle images with an accuracy of 91.4% and 85.6% respectively (Supplementary Fig. 3c). All labels were manually assessed, and subsequently corrected by experts (Supplementary Fig. 3d). Finally, using two orthogonal methods we conducted performance measurement for object classification. First, by using a correlation matrix to compare automated *versus* human segmentation outcomes we found a high correlation coefficient denoting a strong match for both spindle and chromosome categories (Supplementary Fig. 3e). Second, using *apriori* information of the orientation of the spindle pole-to-pole axis and metaphase chromosome plate axis, we confirmed the extent of perpendicular properties between the two automatically annotated objects (mitotic spindle and chromosome plate) in 88 time-lapse movies from at least six independent repeats (Supplementary Fig. 3f). In summary, the manual and computational evaluation efforts together demonstrate a high accuracy with which automated labels are generated using SpinX.

### Benchmarking and refining annotation of classes improved SpinX AI performance

To perform the segmentation of label-free cell cortex and fluorescently-labelled mitotic spindle, we trained and compared two groups of neural network models referred to as ‘SpinX-base’ and ‘SpinX-optimised’. The two models differed in annotation quality and number of epochs, with an increased number for the optimised model (Supplementary Table 1). Annotations for the base model were created by beginner users (0-2 years experience in Cell Biology, N = 800 cortex and 900 spindle images), whereas annotations for the optimised model were created by expert users (>3 years experience in Cell Biology, N = 1300 cortex and 1390 spindle images). For both SpinX-base and SpinX-optimised, data augmentation techniques were carefully selected to artificially increase image variety. For label-free cell cortex images, augmentation was achieved by blurring through Gaussian filtering and contrast normalisation (Supplementary Fig. 4a). This was done to improve the robustness of the cell cortex model in segmenting uniform pixel signals within the cytoplasm. Translation, rescaling, rotation and shearing were added to address the natural variation in cell shape and size (Supplementary Fig. 4a). For mitotic spindle images, higher priority was given to image flipping and rotation in order to better emulate spindle dynamics (Supplementary Fig. 4b).

Model performance was examined through metrics, such as mean Intersection over Union (IoU), mean average precision (AP) and Loss function, allowing the evaluation of classification and mask accuracy (Supplementary Figs. 5 and 6). Comparison of the SpinX-base and SpinX-optimised spindle models suggested an improvement in the ability of SpinX-optimised to generalise, leading to more accurate predictions on unseen data: the AP score for the validation dataset was greater (0.053) in the optimised version, along with marginal improvements in AP (0.012) and Loss reduction (0.0365) for the training dataset (Supplementary Figs. 5, 6 and Table 2). For the cell cortex model, SpinX-optimised displayed a notably higher mean IoU (0.122) and AP (0.124) than SpinX-base for the validation dataset, suggesting the occurrence of less errors during classification and higher accuracy when predicting segmentation masks (Supplementary Figs. 5, 6 and Table 2). To gain further insight on how annotation quality affected model performance, we manually examined the annotations of SpinX-base. 44% and 31% of images required reannotation to precisely outline the boundaries of the cell cortex and spindle, respectively (Table 3). Considering the elevated performance of SpinX-optimised compared to SpinX-base, we conclude that annotation quality and the hyperparameters used to train a Mask R-CNN-based model can greatly affect performance, and so we use SpinX-optimised for subsequent studies.

Next we benchmarked the ability of SpinX to perform segmentation through temporally and spatially discontinuous image sequences. For this, we randomly selected a separate set of 10 time-lapse movies and examined model performance on the cell cortex and the spindle. As routinely performed in the DL field^5,37,44^, we computationally evaluated model performance on spindle and cell cortex segmentation (n = 630 previously unseen images for each model) without (‘native’ Mask R-CNN) and with (SpinX-optimised) post-processing (Fig. 2a and b). We also compared the model performance of SpinX-optimised with U-Net, a DL architecture that has been previously used for cell segmentation^5,7,45^. We trained the U-net based models using the same training datasets (n = 1300 cell cortex model; n = 1390 spindle model) (Fig. 2a). Model performance was evaluated by matching the predictions of each model to the ground truth masks through the IoU metric (Fig. 2a and b). For both spindle and cell cortex segmentation, SpinX outperformed the other two state-of-the-art methods (Fig. 2a).

**Figure 2.**
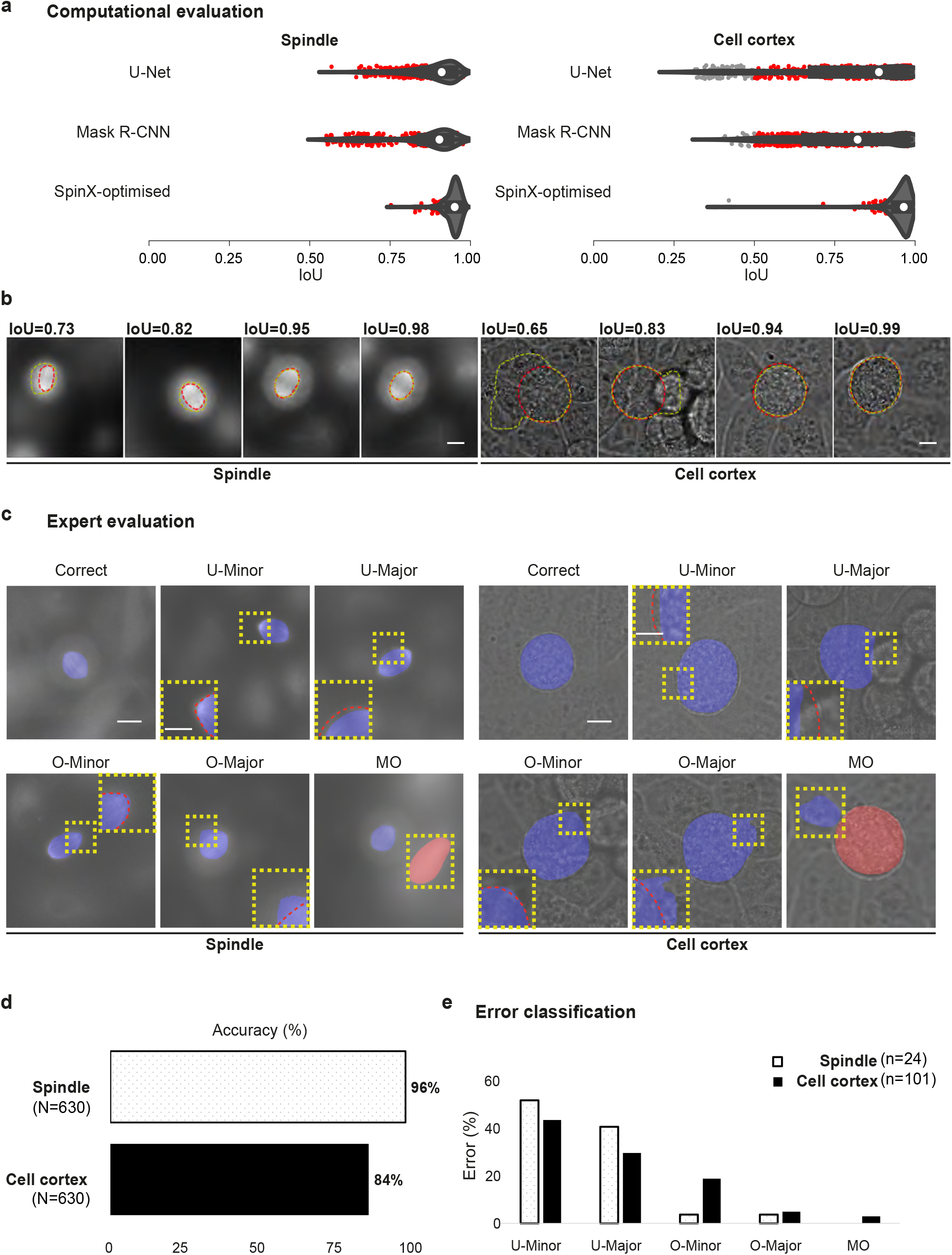
Computational and manual evaluation of SpinX shows high accuracy for spindle and cell cortex segmentation. **a**. Violin plots show the distribution of Intersection over Union (IoU) scores calculated from predictions with U-Net, Mask R-CNN and SpinX-optimised. White marker within the box refers to the median, the shaded area refers to the estimated kernel probability density and the box indicates the interquartile range of the data. Grey and red dots correspond to IoU scores smaller or greater than 0.5, respectively. **b**. Representative images show a range of different IoU scores calculated between the ground truth (red line) and predicted mask (yellow line) for spindle (left) and cell cortex (right). Scale bars: 10 *µ*m. **c**. Representative SpinX prediction images for spindle (left) and cell cortex (right) describing the manual error classification system. Incorrectly segmented images were classified into ‘under segmentation minor’ (U-Minor), ‘under segmentation major’ (U-Major), ‘over segmentation minor’ (O-Minor), ‘over segmentation major’ (O-Major) and ‘multiple objects with artifacts’ (MO). Insets show higher magnification of observed errors. The prediction is highlighted by the blue and red overlays with the corresponding ground truth (red dashed line inside the inset image). Scale bars: 10 *µ*m, 5 *µ*m for inset. **d**. Bar chart shows SpinX’s final accuracy, manually evaluated, for the spindle (white) and cell cortex (black) models. **e**. Bar chart shows the proportion of incorrectly segmented images for each error type defined in (c). For (b), (d) and (e), N=1260 images (630 images each for spindle and cell cortex) from 10 3D time-lapse movies across four independent experiments were considered.

As the output of the AI module is directly fed into the 3D modelling module, accurate boundary information is crucial for reliable 3D tracking of objects. Routinely used IoU metrics, although useful for conventional image segmentation purposes, are insufficient for the purpose of spindle tracking because similar IoU scores can reflect different errors in boundary information (Fig. 2b). For example, the consequence of errors in boundary information near spindle poles will be far more severe than around spindle walls; similar exceptions would apply for cell cortex boundaries (Fig. 2b). Hence, to dissect how the model performance metrics (Fig. 2a and b) translate into accurate segmentation of the spindles and the cell cortex, we developed an error classification system (Fig. 2c). We assessed five types of distinct errors: undersegmentation minor (U-Minor), undersegmentation major (U-Major), oversegmentation minor (O-Minor), oversegmentation major (O-Major), and multiple objects (MO) (Fig. 2c). We benchmarked the SpinX-base models on a large dataset of 630 images with stage 3 of SpinX’s architecture activated (Fig. 1b), which significantly increased the overall accuracy by 35% for the cell cortex model, and 15% for the spindle model (Supplementary Fig. 7). We could confirm that the enhanced accuracy was mainly due to the elimination of wrongly predicted images categorised within the ‘MO’ class (Supplementary Fig. 7). Utilising the SpinX-optimised models (for the same set of 630 images) led to an even greater increase in overall accuracy when compared to SpinX-base - 11% for the spindle model and 10% for the cell cortex model, whereby most errors were found under the ‘U-Minor’ class for both models (Fig. 2d and e). In summary, following different optimisations, SpinX’s final accuracy reached 85% for the cell cortex model and 96% for the spindle model (Fig. 2d).

### Generalisation of SpinX to different spindle markers and distinct imaging systems

As neural network models that accurately segment ‘unseen’ types of data signify longevity and wider applicability, we examined the generalisation capacity of the SpinX framework. Our training dataset consisted of spindles labelled using either mCherry-Tubulin or SiR-Tubulin dye, markers of tubulin subunits (Fig. 3a) which are responsible for assembling and disassembling microtubules of the mitotic spindle (reviewed in^16^). To examine the extent to which SpinX can be generalised, we evaluated the accuracy of SpinX in detecting spindles in time-lapse movie datasets where two different fluorescent marker proteins were fused to two distinct microtubule-binding proteins. First we tested image datasets of cells expressing YFP-tagged Astrin, a microtubule-wall binding protein that can be found at the chromosome-microtubule attachment site soon after the tethering of microtubule-ends to chromosomes^46^ (n = 330 images from 10 cells) (Fig. 3b). Model evaluation was carried out by an expert user through manual binary classification of either ‘correct’ or ‘incorrect’ prediction. Expert evaluation showed that SpinX can successfully segment spindles in movies of YFP-Astrin expressing cells with an 88% accuracy (Fig. 3c). The images in the YFP-Astrin dataset were not complete images of the entire mitotic cell but instead cropped images encompassing the spindle structure alone, requiring padding (see Methods) to allow segmentation through SpinX. Next, we tested image datasets of cells expressing mRFP-tagged End-Binding 3 (EB3), a growing microtubule-end binding protein^47^ that can be found at the chromosome-microtubule attachment site and spindle poles^16^ (n = 1540 images from 5 cells) (Fig. 3b). In addition to widefield images, we extended our evaluation to high-resolution confocal images of cells expressing mKate2-EB3 (Fig. 3b; n = 1920 images from 5 cells). Expert evaluation showed that SpinX is equally successful in segmenting spindles of EB3 marker expressing cells in both widefield and confocal microscopy images, with a 95% and 96% accuracy, respectively (Fig. 3c). Thus, the successful segmentation of EB3 and Astrin-labelled spindle datasets demonstrate a striking generalisation capacity of the SpinX framework for a variety of spindle markers and microscopy methods.

**Figure 3.**
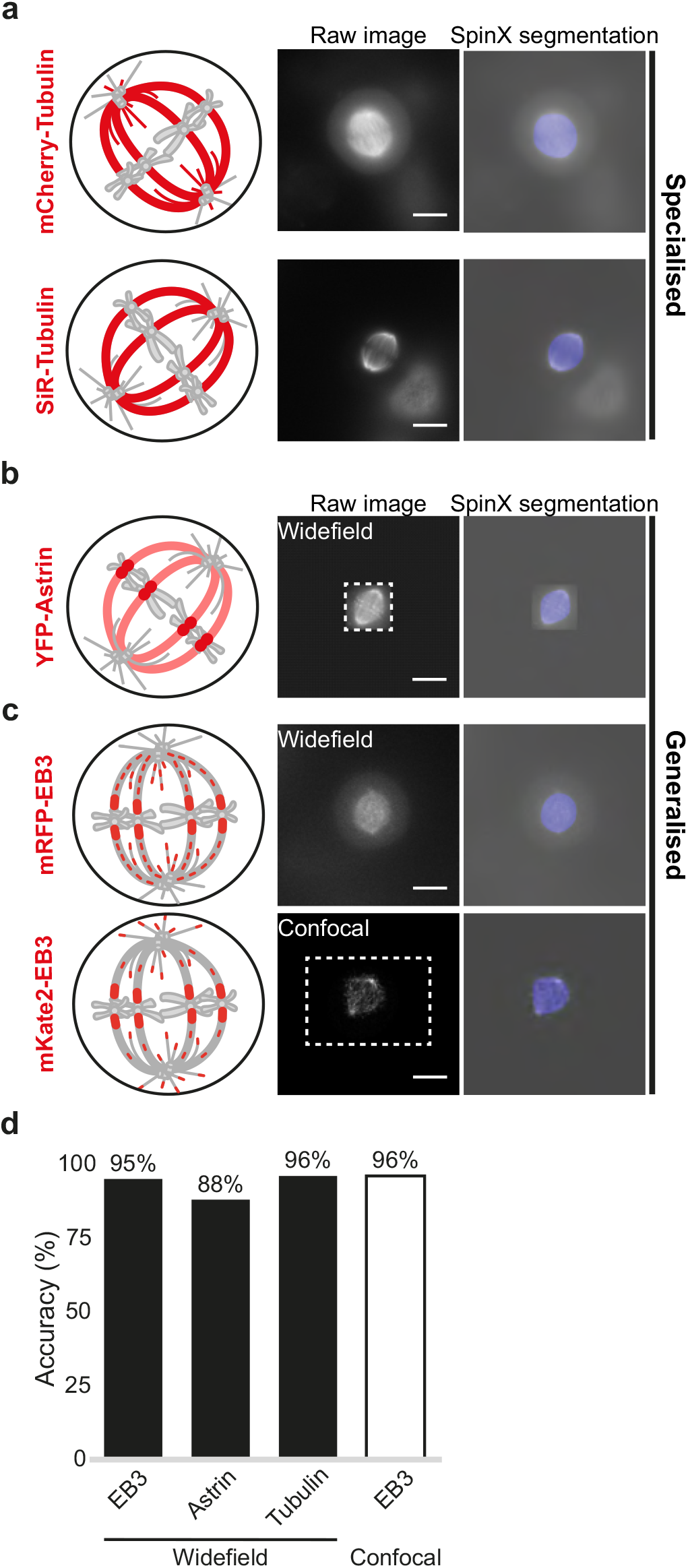
Generalisation of SpinX to segment images of new spindle marker proteins. **a**. Representative images and SpinX segmentation output of mCherry-Tubulin and SiR-Tubulin datasets used to train the spindle model (specialised). **b**. Representative images and SpinX segmentation output of YFP-Astrin, mRFP-EB3 and mKate2-EB3 datasets used to assess the extent to which the spindle model can be generalised. Dashed box shows the original image border (YFP-Astrin and mKate2-EB3 datasets were padded for analysis). Cartoons in **a, b** and **c** show differing localisation patterns (red) of spindle marker proteins. Images acquired using a widefield or a higher resolution confocal microscope are highlighted. **d**. Bar graph shows SpinX’s segmentation accuracy of spindles labelled using mRFP-EB3, YFP-Astrin and Tubulin (mCherry-Tubulin and SiR-Tubulin combined) acquired with a widefield microscope, and mKate2-EB3 using a confocal microscope as indicated. Accuracy was manually scored by experts using the error classification system indicated in Fig. 2c and e. Scale bars: 10 *µ*m.

### Modelling to quantify 3D spindle movements relative to the cell cortex

Reconstructing a 3D spindle structure and cell cortex from 2D slices is a significant challenge in part due to missing information between the *z*-steps. To track spindle movements with reference to the cell cortex, we used the cell cortex prediction mask from SpinX’s AI module (Figs. 1 and 2) to reconstruct the 3D shape of each individual cell (Fig. 4a). Although mitotic cells generally assume a distinct spherical shape^48^, measuring the eccentricity of cell cortex segmentation masks of 96 cells (Supplementary Fig. 8), yielded a median value of 0.3 (a value of 0 being a perfect circle) suggesting that using an ellipsoidal rather than a spherical shape may lead to a more reliable 3D reconstruction.

**Figure 4.**
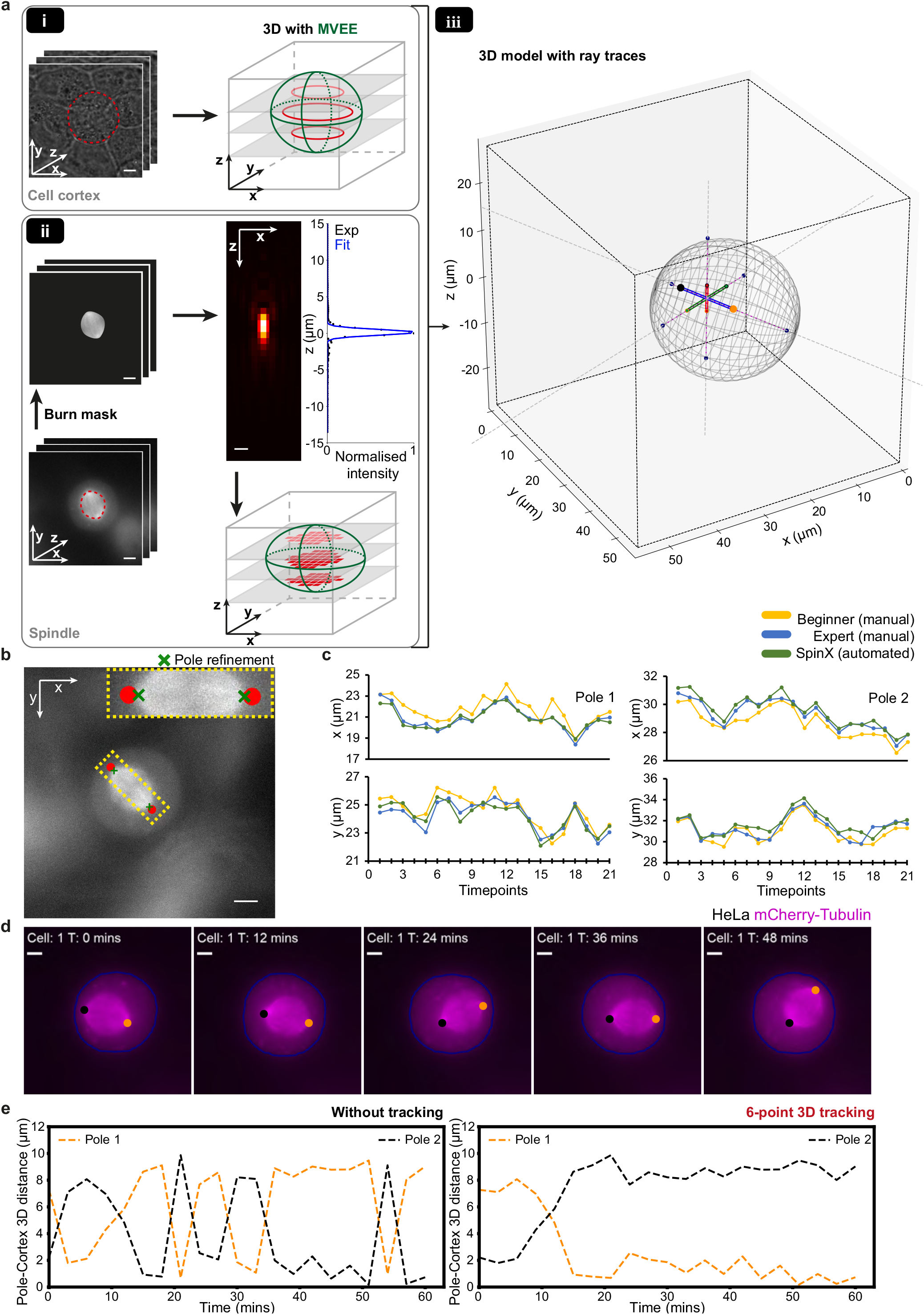
3D reconstruction and modelling for time-resolved analysis of spindle-cortex interaction changes. **a**. (i) Representative time-lapse images of a mitotic cell (left) with the corresponding outlined masks (red dashed line) predicted with SpinX’s AI module. SpinX utilises *z*-slices of cell cortex boundaries (red) to reconstruct the 3D shape of the cell via Minimum Volume Enclosing Ellipsoidal fit, MVEE (green). (ii) Representative time-lapse images of a spindle with the corresponding outlined masks (red dashed line) predicted with SpinX’s AI module. Merging masks with raw images (burn) removes non-spindle signals. PSF was generated and fitted (blue line) to map intensity fluctuations with changes in axial positions for each pixel (red) belonging to the spindle. To reconstruct the 3D structure of the spindle, MVEE (green) was applied. (iii) The 3D plot shows the complete model. The cell cortex is represented by the polygon mesh in grey with the spindle principal axes which corresponds to spindle height (red), width (green) and length (blue). The large orange and black dots at the ends of the length axis correspond to the spindle’s individual poles, while the smaller dots correspond to the ends of the spindle height (red-filled) and width (green-filled) axes. Ray traces from the spindle poles are represented by dashed grey lines and their intersection points are marked as dark blue dots on the cortical mesh. The pole-cortex distance is outlined in magenta. **b**. Demonstration of pole refinement in SpinX. Spindle pole estimations without and with pole refinements are indicated by red dots and green crosses, respectively. **c**. Representative line plots show x and y coordinate changes of a spindle tracked through time, for its poles 1 and 2, measured either manually by a beginner (yellow), expert (blue) or automatically with SpinX (green). **d**. Representative max projection time-lapse images of a HeLa cell expressing mCherry-Tubulin. Orange and black dots represent spindle poles 1 and 2, respectively. Cell outline (blue) was extracted from the predicted segmentation mask of cell membrane by SpinX’s AI module. **e**. Line plots show individual pole-cortex 3D distance measurements computed from (d) for pole 1 (orange) and pole 2 (black) without and with 3D 6-point tracking, respectively. Scale bars: 5 *µ*m and 1 *µ*m for PSF in (b).

We reconstructed the 3D structure of the cell cortex using label-free brightfield images by applying a Minimum Volume Enclosing Ellipsoid fit (MVEE) on the boundary pixel coordinates extracted from the prediction mask of the cell cortex (Fig. 4a-i). To reconstruct the 3D structure of the spindle using fluorescent images, we took advantage of the point-spread function (PSF) of our imaging system. The PSF enabled the estimation of the *z*-coordinates of spindle-associated pixels, which were subsequently used to reconstruct the spindle’s 3D structure by applying the MVEE (Fig. 4a-ii). To capture spindle movement relative to the cell, we measured spindle pole-to-cortex distances (Fig. 4a-iii purple line). For this, we modelled 3D ray traces where we analytically identify the intersection points between the spindle’s principal (pole-to-pole) axes and the rounded cell cortex. Thus the line that passes the two intersection points (Fig. 4a-iii dark blue dots) at the cortex, will pass through the spindle axis of interest as well (see Methods; Fig. 4a-iii dashed grey and purple line; and Supplementary Fig. 11 purple line). This required that the spindle poles are precisely identified, hence, we bench-marked the extent to which MVEE could accurately identify spindle length (the long-axis of the ellipsoid). We observed that due to the intrinsic structure of the spindle, MVEE tends to overestimate the spindle length which in turn alters the predicted spindle pole position (Fig. 4b). This bias accounted for a median spindle pole displacement of 0.5-1 *µ*m in SpinX, compared to manual analysis, which is a 5-6% difference in total spindle displacement (N=4 cells, 84 instances) (Supplementary Fig. 9). This could be recalculated by extracting 3D coordinates along the spindle’s pole-to-pole axis to identify the first and last occurrence of high intensity values that were then assigned as refined spindle pole locations (Table 1). Comparing spindle pole locations obtained either with SpinX or manually (both beginners and experts) confirmed that SpinX’s measurements with the pole refinement algorithm closely matches with the positions tracked manually by an expert, while outperforming manual assessment by a beginner (Fig. 4c).

**Table 1.**
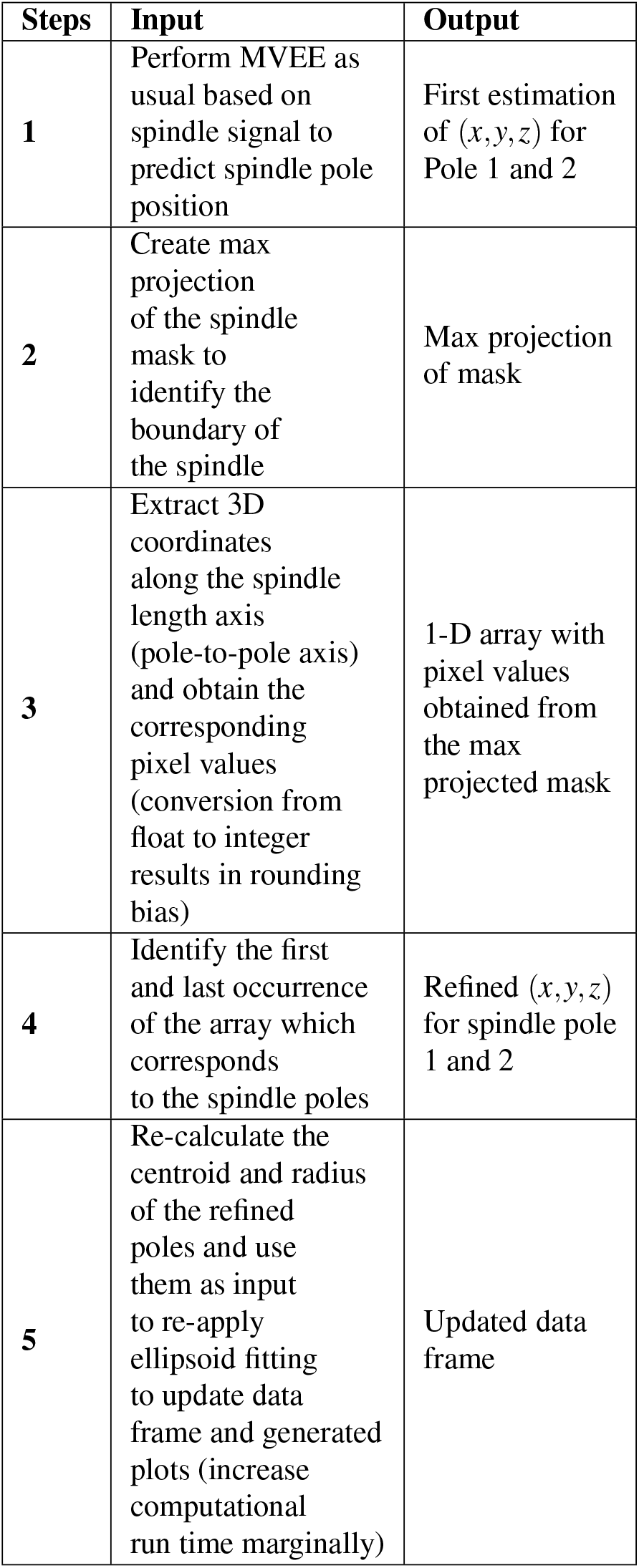
SpinX pole refinement algorithm to compensate overestimation of spindle length axis.

| Steps | Input | Output |
| --- | --- | --- |
| 1 | Perform MVEE as usual based on spindle signal to predict spindle pole position | First estimation of $(x, y, z)$ for Pole 1 and 2 |
| 2 | Create max projection of the spindle mask to identify the boundary of the spindle | Max projection of mask |
| 3 | Extract 3D coordinates along the spindle length axis (pole-to-pole axis) and obtain the corresponding pixel values (conversion from float to integer results in rounding bias) | 1-D array with pixel values obtained from the max projected mask |
| 4 | Identify the first and last occurrence of the array which corresponds to the spindle poles | Refined $(x, y, z)$ for spindle pole 1 and 2 |
| 5 | Re-calculate the centroid and radius of the refined poles and use them as input to re-apply ellipsoid fitting to update data frame and generated plots (increase computational run time marginally) | Updated data frame |

### Tracking of 3D spindle movements through time

To study temporal changes in the spindle’s 3D position, we implemented a six-point tracking algorithm based on the *k*-NN algorithm. The six points represent the end points of the three principal axes of the ellipsoid which correspond to spindle height, width and length axes. By assigning the smallest euclidean distance to pole-1 and not pole-2, we ensured the correction of falsely assigned pole identities through time (see Methods). To test how frequently corrections have to be assigned, we analysed 10 randomly selected time-lapse movies. Correction with *k*-NN was required for around half of the time for spindle width and length axes, and one third for spindle height axis (Supplementary Table 4). To evaluate the impact with and without tracking, we measured the 3D distances from each spindle axis to the cell cortex (Fig. 4d). The pole-to-cell cortex 3D distance was obtained by computing ray traces. We confirmed that consistent pole assignments with the tracking algorithm enabled accurate measurements of spindle pole positions through time (Fig. 4e). Thus, changes in spindle pole-to-cell cortex distances, as a measure of spindle displacement, could be tracked in 3D through time (Supplementary Video 1)

### SpinX reveals *in vivo* use of MARK2 kinase inhibitor

To showcase the strength of an AI-based spindle tracking tool for analysing the movements of 100s of spindles through time, we set out to quantify perturbations in spindle movements in the presence of an inhibitor of MARK2 (Microtubule affinity regulating kinase 2, Par1 kinase family), implicated in centering spindles along the equatorial axis^25^. Although a screen for drugs with therapeutic potential had identified an *in vitro* inhibitor of MARK2/Par1b activity (hereafter: MARK2i; Calbiochem 39621)^49^, whether this inhibitor can rapidly disrupt MARK2’s function in spindle centering is not known. We first collated 3D time-lapse movies of HeLa cells expressing H2B-GFP and mCherry-Tubulin (Fig. 5a) exposed to MG132 (to enrich them in metaphase), whereby they were imaged in the presence or absence of MARK2i for up to 3 hours. Visual inspection of time-lapse movies suggested that spindles of MARK2i treated cells may be equatorially off-centered in some but not all time-frames (Fig. 5b). To quantitatively assess changes in spindle movement in 3D, we used SpinX for tracking and decomposing spindle movements in longitudinal, equatorial and axial orientations with respect to spindle length, width and height axes, respectively (Supplementary Fig. 10). By including the 3D cell cortex information, we could additionally account for variability in cell-to-cell differences i.e., the available space for spindles to move. In control metaphase cells (DMSO treated cells) - as expected^27^ - we observed a bias towards longitudinal movements of the spindle along the pole-to-pole axis and highly restricted equatorial movements (Fig. 5c). However, in MARK2i treated cells (N=12), the fraction of longitudinal movement is significantly reduced, while the fraction of equatorial and axial movement are both increased compared to control cells (N=11) (Fig. 5c). The strong increase in equatorial movement following MARK2i treatment shows that MARK2 inhibition promotes equatorial movement, similar to MARK2 depletion^25^, revealing an immediate *in vivo* impact of the inhibitor.

**Figure 5.**
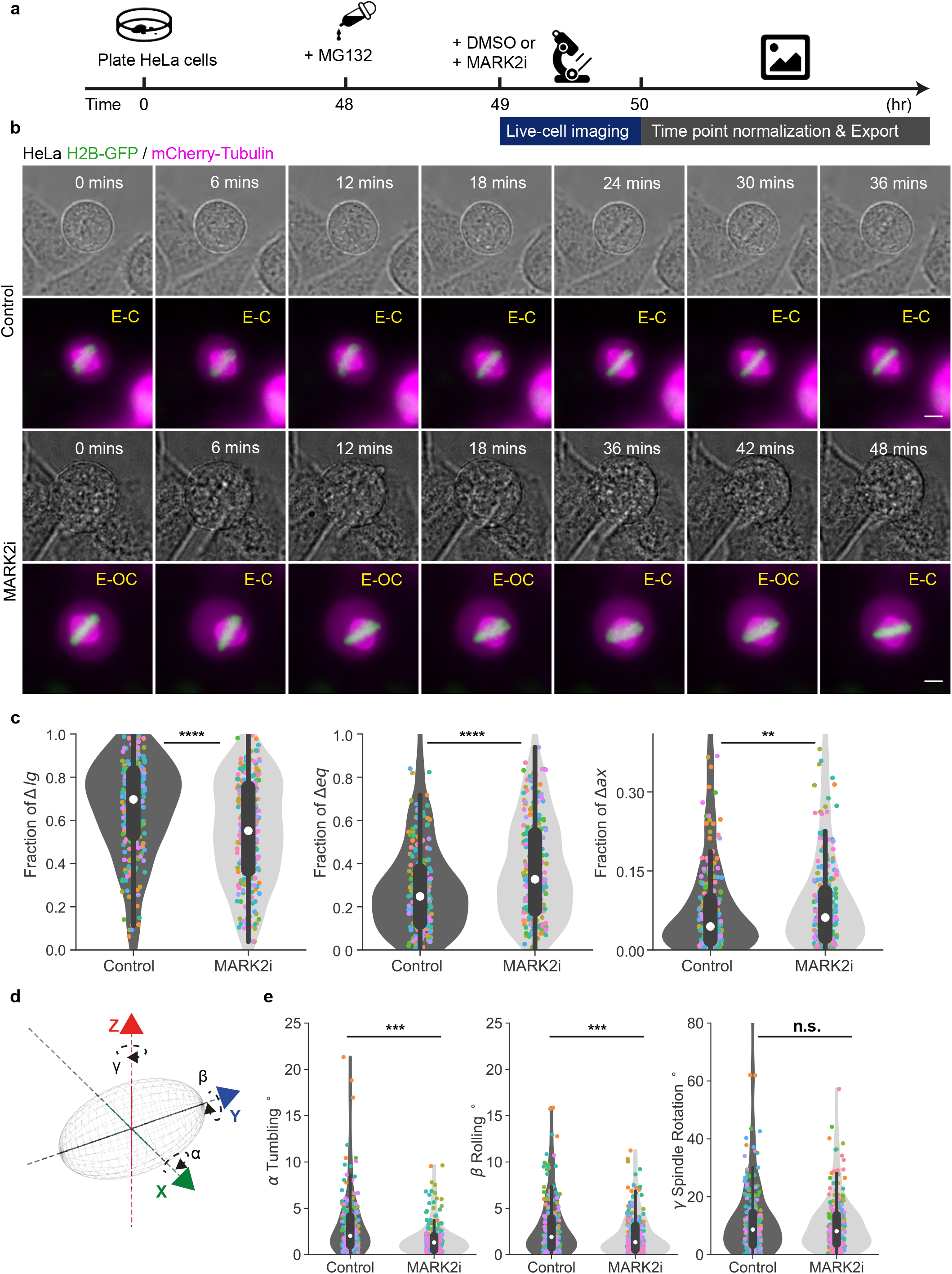
MARK2i promotes equatorial movement of the mitotic spindle. **a**. Experimental regime. HeLa cells were exposed to MG132 (10 *µ*M) 1 hour before imaging. Either DMSO (control) or 5 *µ*M MARK2 inhibitor (MARK2i) as indicated were added during imaging. **b**. Representative brightfield images displaying cell cortex (grey) and maximum projection live-cell images of a HeLa cell expressing H2B-GFP (green) and mCherry-Tubulin (magenta). Equatorial centered and off-centered spindles are marked (E-C) and (E-OC), respectively. Cells were imaged over 1 hour with images taken every 3 minutes. **c**. Violin plots show fractions of longitudinal (Δlg), equatorial (Δeq) and axial (Δax) spindle displacement. Corresponding coloured dots represent measurements from all time points of the same cell. White marker within the box refers to the median, the shaded area refers to the estimated kernel probability density and the box indicates the interquartile range of the data. **d**. Cartoon shows a 3D spindle (grey) with the corresponding rotation angles *α* (spindle tumbling), *β* (spindle rolling) and *γ* (spindle rotation) along its principal spindle axes *x, y, z*, respectively. **e**. Violin plots show 3D angle distribution for *α* spindle tumbling, *β* rolling and *γ* rotation. Corresponding coloured dots represent measurements from all time points of the same cell. White marker within the box refers to the median, the shaded area refers to the estimated kernel probability density and the box indicates the interquartile range of the data. Statistical significance was determined by Mann-Whitney U test (in c and e) after a pre-analysis of the underlying distribution with a Shapiro-Wilk test. N=11 control and N=12 MARK2i cells across three separate experiments. Scale bars: 5 *µ*m.

A defect in anaphase spindle orientation along the interphase long-axis after MARK2 depletion has been reported^25^, but changes in metaphase spindle orientation have not been previously quantified. We took advantage of SpinX’s reconstructed spindle principle axis and its corresponding centroid to compute 3D rotational angle changes in *α* spindle tumbling, *β* rolling and *γ* rotation in metaphase spindles of control and MARK2i treated cells (Fig. 5d and e). We find that the extent of spindle rotation is not affected upon MARK2i (Fig. 5e), but both spindle tumbling and rolling movements are significantly reduced. To test if this reduction is time-dependent, we performed correlation analysis between angle changes and time (Supplementary Fig. 10). Computed Pearson correlation coefficients *ρ* showed no linear correlation in both conditions in spindle tumbling (*ρ* = 0.101 and *ρ* = 0.060), spindle rolling (*ρ* = 0.102 and *ρ* = −0.090) and spindle rotation (*ρ* = 0.030 and *ρ* = 0.051). These findings reveal that MARK2i treatment alters spindle tumbling and rolling movements, but not rotational movements.

As SpinX-based spindle tracking helped uncover the immediate *in vivo* impact of the MARK2 inhibitor in mitotic cells for the first time, we used the same concentration of MARK2i that altered spindle movements (Fig. 5) to test if transient exposure to the inhibitor is sufficient to alter MARK2 localisation during interphase. While MARK2 wildtype (WT) localises as puncta throughout the interphase cell substrate, the kinase dead (KD) mutant localises as long striations parallel to actin fibres^50^. Following a brief 30-minute exposure to MARK2i, MARK2-YFP localised as long striations parallel to actin fibres near the cell substrate (Supplementary Fig. 12a and b). In contrast to prominent punctate-foci distribution of MARK2-YFP in control cells treated with DMSO, MARK2i treated cells showed fewer punctate-foci but a higher number of long striations along the actin stress fibres (Supplementary Fig. 12b). Segmentation and quantification of eccentricity of MARK2-YFP foci (Supplementary Fig. 12c) showed that the localisation of MARK2-YFP was significantly altered upon MARK2 inhibition, representing eccentricity values similar to the distribution of foci in MARK2-KD expressing cells (Supplementary Fig. 12d). Prolonged MARK2 inhibitor treatment, by exposing cells for a longer period (16 hours), did not significantly change the localisation pattern compared to a shorter period of drug treatment (Supplementary Fig. 12d), demonstrating the *in vivo* use of the inhibitor to acutely block MARK2 function during both interphase and mitosis. Thus, SpinX enabled precise tracking of 3D spindle movements following inhibitor treatment, showcasing the significance of DL-based quantitative analysis of discontinuous time-lapse movies.

## Discussion

We show that an AI-based image analysis framework supported by 3D modelling can better harness dynamic information in time-lapse microscopy movies. By bringing together large-scale time-lapse movie datasets and a computational framework, SpinX, we can now precisely track spindle movements in 3D using diverse spindle protein markers. Using manual and automated benchmarking tools we establish that SpinX can reliably (i) detect and segment the spindle and the cell membrane, (ii) transform 2.5D data to true 3D through ellipsoid reconstruction and (iii) track spindle movement relative to cell size through 3D mathematical modelling. The methods we present here can be of general use beyond spindle tracking: 3D reconstruction for fluorescent images by utilising properties of the PSF, ray-tracing principle to model 3D movements relative to different subcellular structures, a 6-point 3D tracking algorithm for capturing translational and rotational movements of structures, and an expert error classification system to support model evaluation and refinement. Lastly, we showcase SpinX’s potential in supporting preclinical and drug development studies by taking advantage of the sensitivity of spindle movements to subtle molecular or subcellular changes following transient exposure to a chemical inhibitor of MARK2 kinase.

One of the major hurdles in DL-based tool development is the lack of large volumes of high-resolution time-lapse datasets that are essential for feature-rich analysis of subcellular structures. However, the lack of sophisticated image analysis tools discourages the generation of large-scale high-resolution datasets. Here we break this conflicting ‘catch-22’ loop by generating both the time-lapse movie dataset and analysis tools for measuring and characterising spindle size, position and movements. Thus, SpinX provides a complete framework including modules for annotation, training, modelling and analysis, and the possibility of validating predictions at multiple steps of the process (Fig. 6), enabling robust 3D tracking of spindle movements relative to the cell cortex.

**Figure 6.**
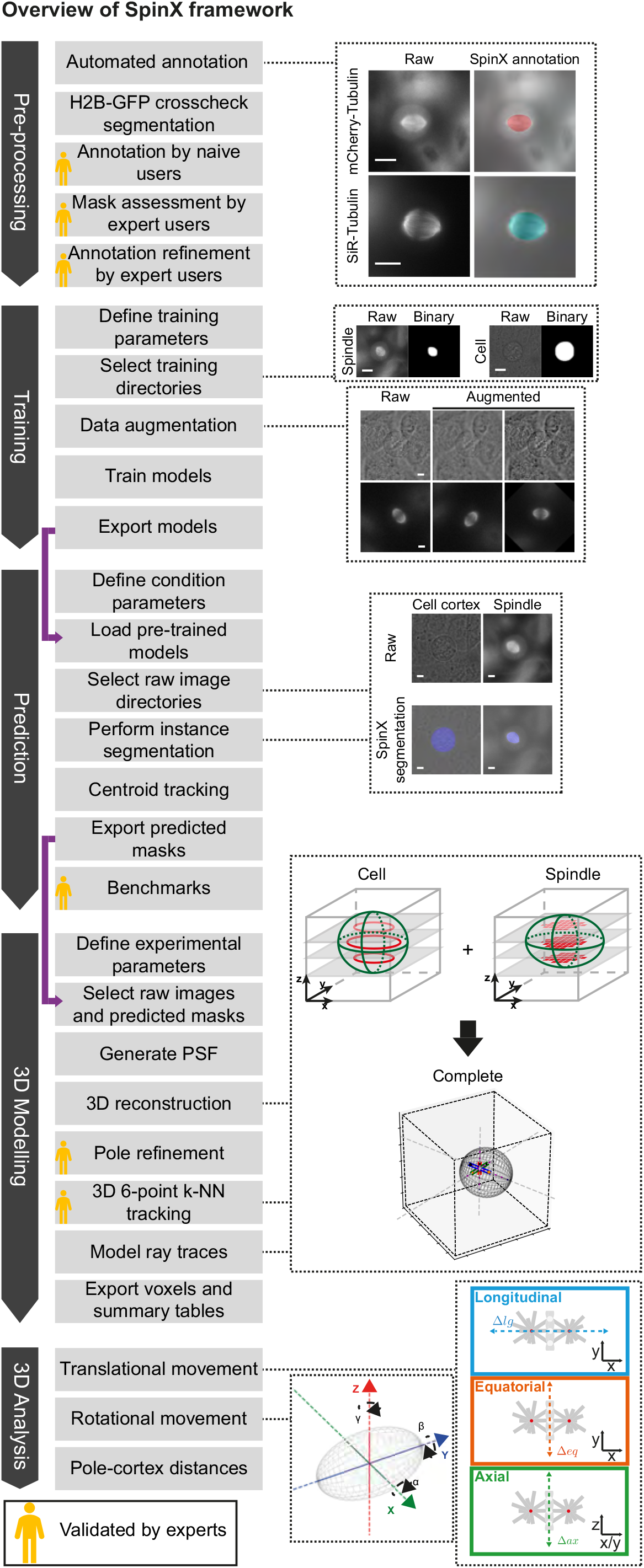
SpinX’s comprehensive framework. Diagram shows the complete framework of SpinX including modules for generating annotations (pre-processing), training, prediction, 3D modelling and 3D analysis (dark grey arrows). Each module has a series of automated and manual steps (light grey boxes), with purple arrows indicating how data is passed between modules. Representative raw images and their corresponding automatically annotated spindle images, along with raw and binary spindle and cell cortex images belonging to the training dataset are shown. 3D reconstructions of the cell cortex and mitotic spindle enable the extraction of translational and rotational spindle movements, and pole-to-cortex distances. Image annotation, predicted masks enabling temporal and spatial links between images, and 3D modelling of pole position and tracking are all validated by experts (gold icons). Scale bars: 10 *µ*m.

SpinX’s contribution for the live-cell microscopy field is 2-fold: extending the Mask R-CNN network to perform predictions on 3D time-lapse movie datasets, and a high-resolution annotated dataset of images of fluorescent labelled spindles and label-free cells. The Mask R-CNN-based architecture allowed us to harness the network’s flexibility in handling images of an arbitrary size, and supported instance segmentation of multiple classes - a crucial feature for cell segmentation to separate overlapping cells, while also classifying them into distinct phenotypes and providing unique IDs. The evaluation of the neural network model using a detailed error classification system helped assess the strengths and limitations of SpinX. For example, errors in cell cortex prediction were mostly categorised as ‘undersegmentation’ which were mathematically compensated by the MVEE during modelling. Our findings highlight the importance of annotation quality, especially for studies where precise measurements of object boundaries are important for accurate 3D-modelling and object tracking through time. Benchmarking studies with expert and beginner users confirmed the benefits of Mask R-CNN, including the generalisation capacity of SpinX to detect spindle markers beyond the ones used to train the model.

Unlike manual analysis of spindle movements, SpinX can capture translational and rotational movements of subcellular structures using the 6-point 3D tracking algorithm, and measure 3D spindle movements relative to the cell cortex using principles from ray tracing methods. These allowed the first careful assessment of the impact of the MARK2 inhibitor on spindle movements. MARK2 inhibitor treatment does not affect spindle rotation per se, but affects spindle rolling and tumbling by altering the equatorial positioning of spindles. These findings are consistent with equatorial positioning defects, previously reported through manual analysis of MARK2 depleted cells^25^.

As the mitotic spindle movements are highly sensitive to changes in the cells’ cytoskeleton, membrane compartment and chromosome position, SpinX is expected to help accelerate and advance automated screening of drug targets and chemical compounds. SpinX developed for single cell studies, based on Mask R-CNN, can be readily generalised to multi-cell images and also multi-content images to allow the simultaneous tracking of more than one subcellular structure.

## Methods

The SpinX framework was developed in Python 3, using numpy, scipy, tensorflow, keras, scikit, pandas and opencv. For the interactive interface tkinter was used. Figures were generated using matplotlib, seaborn and jupyter-notebook.

### Data generation

#### Cell line and culture conditions

HeLa cell lines used were cultured in Dulbecco’s Modified Eagle’s Medium (DMEM) supplemented with 10% fetal bovine serum (FBS; ThermoFisher 10270106), 1% Penicillin/Streptomycin (ThermoFisher 15140122) and 0.1% Amphotericin B (Fungizone; ThermoFisher 11510496). Cell lines were cultured as a monolayer at 37°C and 5% CO_2_. HeLa H2B-GFP, mCherry-Tubulin cell line was generated by transfecting mCherry-Tubulin expressing eukaryotic plasmid vector into HeLa H2B-GFP cells^51^. The HeLa mKate2-EB3 cell line was generated by transfecting a pmKate2-EB3 plasmid vector (Evrogen #FP316) into HeLa cells. Plasmid transfection was carried out using DharmaFECT duo (Dharmacon T-2010). The HeLa H2B-GFP, SiR-Tubulin cell line was generated by adding SiR-Tubulin dye, a paclitaxel-based fluorescent compound^52^ (Spirochrome SC002; 100 nM) just 1 h before imaging. The HeLa FRT/TO cell line expressing siRNA-resistant MARK2-YFP-WT or KD mutant was generated by transfecting a Tet-inducible expression vector encoding siRNA-resistant MARK2-YFP-WT or KD, followed by colony picking. Vectors bearing point mutants of MARK2 were generated by polymerase chain reaction-based point mutagenesis and confirmed by DNA sequencing^50^. Fluorescent cells were enriched using a BD FACSAria III Cell Sorter for fluorescence-activated cell sorting (FACS).

#### Live-cell microscopy

Live-cell imaging experiments were performed using cover glass chambered dishes (ThermoFisher 155383PK). MG132 (Tocris Biosciences 1748; 10 *µ*M) was added 1 h before imaging to synchronise cells at metaphase^39^. During imaging, cells were incubated in Leibovitz’s L15 medium (ThermoFisher 11415064). For MARK2 studies, MARK/Par-1 activity inhibitor^49^ (MARK2i; Calbiochem 39621; 5 *µ*M or 10 *µ*M) was added prior to imaging. For HeLa FRT/TO MARK2-YFP (WT and KD) experiments, Doxycycline (Fisher Scientific 10224633; 200 ng/ml) was added 16 h prior to imaging. SiR-Actin dye^52^ (Spirochrome SC001; 100 nM) was added 30 minutes before imaging. All imaging sessions were conducted in a chamber at 37°C.

Widefield images were acquired with an Applied Precision Deltavision Core deconvolution microscope equipped with a dual camera system composed of a CoolSNAP and Cascade2 Camera (Photometrics) under EM mode. For live-cell studies, images were taken every 3 minutes (21 timepoints - total time 60 mins) with optimised exposure times ranging from 0.1-0.2 seconds depending on the imaging channel. For each experiment, at least 3 *z*-sections (2 *µ*m gap) were acquired using an oil-based 60X NA 1.42 objective or 100X NA 1.40 objective. High-resolution imaging datasets have pixel sizes ranging between 0.04144-0.06887 *µ*m. Time-point equalisation, deconvolution and data export (Tiff-format) were performed in SoftworX 6.5.2.

Confocal images were acquired using a Leica Stellaris 8 confocal microscope with an oil-based 63X NA 1.40 objective. Each movie consisted of at least 4 *z*-sections (max 46) taken with 0.2-0.5 *µ*m gaps. All movies underwent adaptive deconvolution (Lightning mode). Before processing through SpinX, movies were converted to 8-bit and padded to 1024×1024 dimensions using our padding algorithm (described below).

#### Image Datasets

Our image pools include 13,230 cell membrane and 15,120 spindle images. Cell membrane images were pooled from 188 3D high-resolution live-cell movies, while the spindle images were pooled from 217 3D high-resolution live-cell movies, both across 26 experiments. A uniform random generator was used to randomly select 2180 cell membrane and 2320 spindle images to build the training, validation and testing datasets (Supplementary Fig. 1). For SpinX’s final cell membrane model (i.e. SpinX-optimised) the training dataset consisted of 1300 images (Tables 1, 2, 3, and Supplementary Figs. 5 and 6), the validation dataset consisted of 250 images (Tables 3, and Supplementary Figs. 5 and 6) and the testing dataset consisted of 630 images (Fig. 2). For SpinX’s final spindle model (i.e. SpinX-optimised) our training dataset consisted of 1390 images (Tables 1, 2, 3, and Supplementary Figs. 5 and 6), the validation dataset consisted of 300 images (Tables 2, 3, and Supplementary Figs. 5 and 6) and the testing dataset consisted of 630 images (Fig. 2). For testing the generalisation extent of SpinX high-resolution 3D live-cell time-lapse datasets of HeLa cells expressing mRFP-EB3 (1540 images, from 5 cells), YFP-Astrin (330 images, from 10 cells), and mKate2-EB3 (1920 images, from 5 cells) were used.

#### Annotation

Manual annotations required for training and evaluation (i.e. ground-truth masks) were performed with VGG Image Annotator (VIA) tool^53^. Any missegmented images from SpinX’s AI output were also manually corrected through VIA before 3D reconstruction and modelling. Automated annotations were generated through chromosome and spindle segmentation pipelines combining different conventional image processing techniques. The chromosome segmentation pipeline used for segmenting H2B-GFP labelled chromosomes includes: (1) a median filter for noise reduction^54^; (2) Otsu’s method for iterative two-class thresholding (by minimising the weighted within-class variance), thereby globally reducing a greyscale image to a binary image^55^; (3) a connectivity matrix making up an 8-connected neighborhood used for clearing any pixels found at the image border; and (4) contour smoothing with the Savitzky-Golay signal processing filter^56^ (Supplementary Fig. 2a). The spindle segmentation pipeline used for segmenting SiR-Tubulin labelled spindles includes: (1) median filtering of the size [20, 20] for improving signal-to-noise ratio; (2) an adaptive threshold for estimating the average background illumination intensity; (3) binarisation along with morphological dilation and erosion for removing artifacts; (4) calculating the convex hull of the segmented spindle halves to allow joining; (5) contour smoothing with the Savitzky-Golay signal processing filter^56^; and (6) fitting an ellipse to obtain spindle properties (Supplementary Fig. 2b). The spindle segmentation pipeline used for segmenting mCherry-Tubulin labelled spindles includes: (1) a Gaussian filter for noise reduction; (2) calculation of the image gradient; (3) an automated snake i.e. active contour model^57^, that uses boundary information from the already segmented chromosomes as initial coordinate points; and (4) inversing the snake, thereby propagating from the center of the spindle towards the outer boundary contour to avoid any cytoplasmic noise (Supplementary Fig. 2c). All automatically-generated annotations were manually assessed and corrected if needed using VIA tool^53^.

### Deep neural network

#### ResNet CNN for DL model of spindle and cortex

The ResNet CNN computes full-image feature maps with an increased depth of 101 layers, therefore achieving an elevated semantic value, despite the progressive loss in spatial resolution. The final feature map generated is fed into the RPN, leading to thousands of propositions of where the object of interest is most likely located, termed as regions of interest (ROIs). The presence of an RPN in the architecture allows the detection of individual cells within densely populated images, while also enabling the tracking of the same object across time through the bounding boxes generated. The RPN defines several sets of bounding boxes by a sliding window approach which is based on a computed IoU metric^42^. The sets of bounding boxes then undergo binary classification and regression in a parallel manner, followed by non-maximum suppression for selecting the most accurate non-overlapping bounding boxes^37,58^. The resulting bounding boxes (i.e. anchor boxes) indicating the same ROI are aligned with each other through bilinear interpolation - also known as the ROIAlign layer, which improves pixel accuracy through the refinement of pooling operations (i.e. object extrema)^37^. Subsequently, the FCN allows the simultaneous prediction of the corresponding class and bounding box for each ROI from the detection network, and the generation of the mask within each ROI from the segmentation network.

#### Data augmentation

For both brightfield and fluorescent images used for training the cell cortex and spindle models respectively, augmentation was carried out on every epoch. Augmentation techniques used included image blurring through Gaussian filtering, contrast normalisation, translation, rescaling between 80-120%, rotating up to 180° or shearing by -8 - 8° (Supplementary Fig. 4). Images also underwent flipping, element-wise addition, simple pixel value addition and multiplication, random pixel dropout of up to 10%, gamma adjustment and cropping (Supplementary Fig. 4). For the cell cortex models priority was given to translation, rescaling and shearing to address the natural variation in cell size and shape; whereas for the spindle models priority was given to rotation and flipping to capture the variety in spindle dynamics (Supplementary Fig. 4).

#### Training

For training our Mask R-CNN models we used strategies from ref^59^. The networks were trained for at least 200 epochs (base models) or 500 epochs (optimised models) with stochastic gradient descent at a learning rate of 0.001, a momentum of 0.9, batch size of one image and a weight decay of 0.001 (Supplementary Table 2). The number of anchors for RPN was set to 512. The detection threshold was set at 90%. Models were initiated with COCO pre-trained weights^60^. The best models were selected based on the lowest loss value in the training and validation datasets. To train U-Net, we used a learning rate of 0.00001 with a batch size of 4 and trained for 500 epochs.

#### Metrics

IoU scores were calculated by quantifying the matching between predictions proposed by DL models and their corresponding ground-truth masks. Average Precision (AP) scores to assess class assignment are defined as *AP* = ∑_*n*_(*R*_*n*_ − *R*_*n*−1_)*P*_*n*_, where *P*_*n*_ and *R*_*n*_ are the precision and recall at the *n*th threshold (Supplementary Table 2). Loss functions were determined as described in ref^37^ (Supplementary Fig. 6 and Table 2).

### PSF to estimate spindle pixels in *z*

PSF was simulated with the Gibson-Lanni model^61^ that accounts for different imaging conditions (Table 5). To translate the empirical measurements of the PSF to a mathematical function, the intensity values on the *x, y* and *z*-sections were fitted. Given a 3D image stack of a fluorescence bead where *x* and *y* are intensity values, (*x*_*c*_, *y*_*c*_) is the centroid coordinates of the brightest spot across the *z*-stack, *h* is the height of the Gaussian and *σ*_*x*_ and *σ*_*y*_ are variances in the *x* and *y* directions where (*σ*_*x*_ ≠ *σ*_*y*_). Then, the 2D Gaussian with *k* = 2 is the product function derived from a multivariate Gaussian **X**_∈(*x,y*)_ ∼ *𝒩*_*k*=2_(*µ*, Σ) where 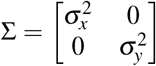 with det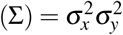:

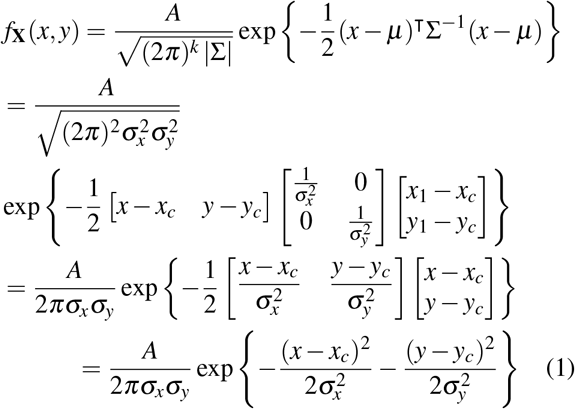

To fit *z* data points of intensity values of the fluorescence bead along the *z*-slices, the Equation 1 can be simplified to a to 1D Gaussian with:

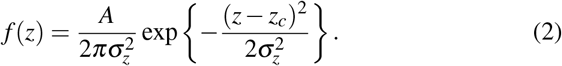

Then, the function that relates the intensity values to the estimated *z*-coordinate 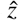 with respect to the reference intensity profile *r*_*int*_ is

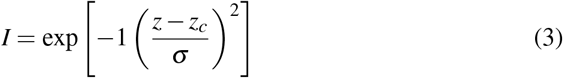

For each pixel, we therefore applied the following equation to estimate its 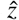-coordinate:

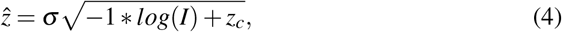

where *z*_*c*_ and *σ* denotes the mean and variance of the Gaussian. To then 3D reconstruct the spindle we use its assigned prediction mask generated by the SpinX AI module and burn it on the initial raw fluorescent image. This step isolates neighbouring signal noise and retains only the pixels belonging to the spindle. Then, we filter the predicted mask by keeping the top 30% of pixels with the highest intensity, thus reducing the number of data points while maintaining the shape of the spindle.

### Ray traces to determine pole-to-cortex distance

Given a line in a three dimensional space which is defined by two points *P*_1_(*x*_1_, *y*_1_, *z*_1_) and *P*_2_(*x*_2_, *y*_2_, *z*_2_), the parametric line for points of intersect can be described by

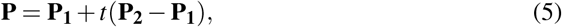

where each coordinate of **P** can be written as

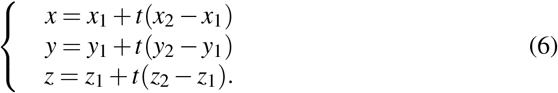

An ellipsoid translated to its center at *P*_3_(*x*_3_, *y*_3_, *z*_3_) can be described by

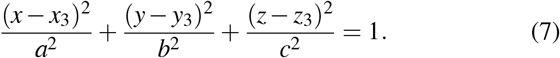

The intersection points *P* of the parametric equation (5) satisfy the substituted equation (6) in (7):

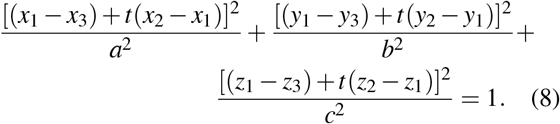

Solving the square values of the parenthesis yields:

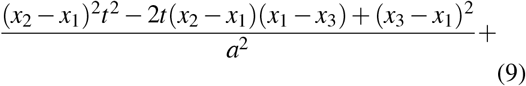

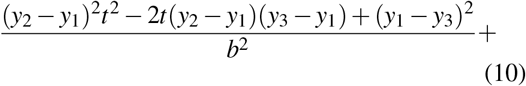

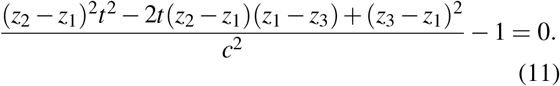

Arranging the expression received as a powers of *t* yields:

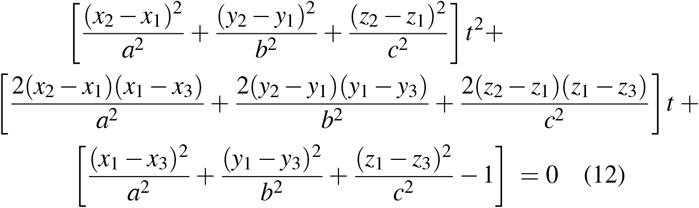

Substituting the equation of the line into the ellipsoid form gives a quadratic equation of the form:

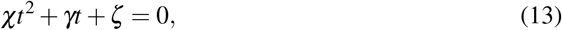

where:

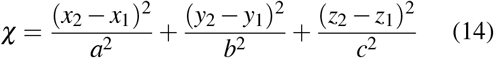

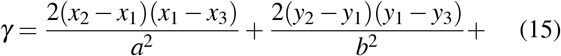

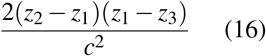

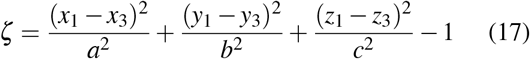

The solution for *t* is then:

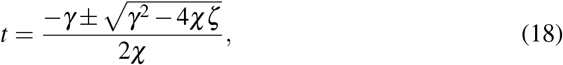

where

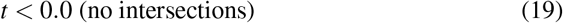

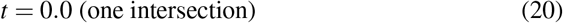

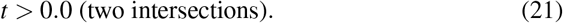

Substituting *t* in (5) yields the intersection points **P** where the spindle axis collides with the cell cortex. By applying the analytical solution, the precise intersection points can be derived at a lower computational cost (Supplementary Fig. 11). Equation 4 was derived by (i) fitting a 2D Gaussian function along the *xy* coordinate of the PSF and (ii) a 1D Gaussian fit at the centroid of the PSF along the *z*-slices.

### SpinX pole location refinement

The spindle pole refinement algorithm takes initial (x,y,z) spindle pole predictions as an input. The spindle boundary coordinates are obtained by taking the maximum projection of the spindle mask. Then, 3D coordinates are extracted along the spindle length axis (pole-to-pole axis) to obtain the corresponding pixel values. The true spindle pole is defined as the first and last occurrence of positive values in the resulting 1D array. Finally, the new position of spindle poles was updated across all data frames for further calculations. Manual analysis used for evaluating the performance of SpinX’s pole position recording was performed on Fiji (ImageJ) software^62^.

### SpinX pole identity assignment

We implemented a six-point tracking algorithm based on *k*-Nearest Neighbour algorithm (*k*-NN). The six points represent the end points of the three principal axes of the ellipsoid which corresponds to spindle height, width and length axis and works as follow:

Given (*x, y, z*) coordinates of *i* = 2 poles at *j* = 2 consecutive time points **P**_1(*t,t* − 1)_ and **P**_2(*t,t* − 1)_, the pairwise distance between poles can be described by the distance matrix

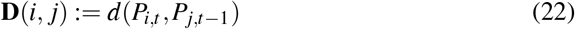

where *d* is the 3D Euclidean distance between two consecutive points

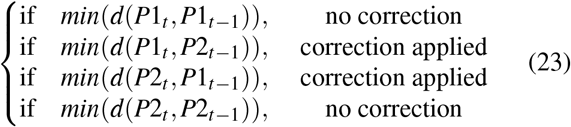

The condition for a correct assignment of pole 1 at *t* is when its distance is smallest at *t −* 1 and largest to pole 2 at *t −* 1. Based on this condition, SpinX perform correction for individual poles whenever they were falsely assigned (e.g. pole 1 at *t −* 1 is closest to pole 2 at *t*). Once corrected, SpinX updates the correction throughout the data frame. To test how frequently corrections have to be applied, we analysed 10 randomly selected time-lapse movies. According to Table 4, correction with *k*-NN was required for around half of the time for spindle width and length axes and one third for spindle height axis, respectively (Fig. 4e).

### Computing 3D rotational movement with Euler’s angle

The extent of spindle rotation is defined by the rotation matrix (**R**_3×3_) which is the product of successive rotation about the *z, y* and *x* axes such as:

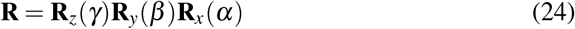

with,

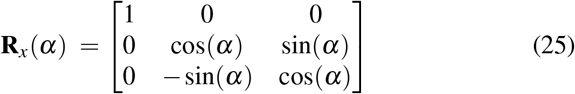

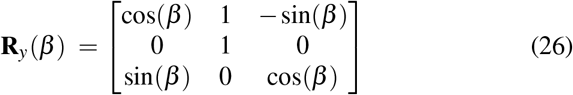

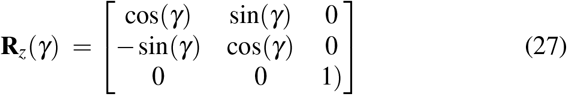

and satisfies:

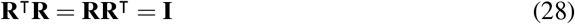

where **I** is the identity matrix.

Then, the corresponding Euler angles *α, β* and *γ* can be computed from the rotation matrix **R** with^63^:

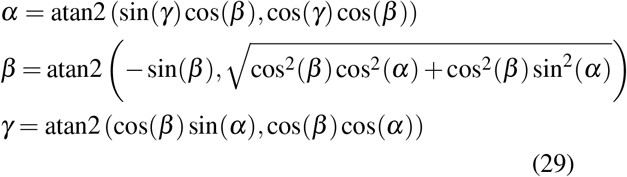

### Padding

Padding of the YFP-Astrin and mKate2-EB3 datasets were performed through an algorithm that extracted small-sized patches of the input image based on their sum of pixel intensity values. The patches exhibiting the lowest sum of intensity values were then used to pad the input image to a desired size. Therefore, the low-intensity small-sized patches emulated and propagated the ‘background’ of the input image. This transformed the Astrin-YFP dataset to 512×512 pixel images and the mKate2-EB3 dataset to 1024×1024 pixel images, subsequently enabling SpinX’s AI module to segment spindles.

### Statistical analysis

Statistical tests were performed in Python 3 (using scipy package) and R-Studio (using R 3.6). Statistical tests were performed on a significance level of *p* ≤ 0.01 or *p* ≤ 0.05. For *p*-values, the following convention holds: not significant (n.s.) with *p >* 0.05, (*) with *p* ≤ 0.05, (**) with *p* ≤ 0.01, (***) with with *p* ≤ 0.001 and (****) with *p* ≤ 0.0001.

## Acknowledgements

We would like to acknowledge funding support from the Biotechnology and Biological Sciences Research Council (BBSRC) to V.M.D ([R01003X/1]), D.D. (LIDo-DTP studentship [BB/M009513/1]) and C.E. (LIDo-iCASE studentship [BB/T008709/1]).

We thank Haajr Faheem (Draviam group), Evangeline Marianesan (Draviam group) and Xinhong Song (Draviam group) for their support in generating pre-annotations; Dr Sagar Joglekar (Sastry group) for his input on the automated annotation pipeline; and Dr Asifa Islam (Draviam group) for providing the YFP-Astrin dataset.

We would like to thank Professor Conrad Bessant (Queen Mary University of London), Dr Chema Martin-Duran (Queen Mary University of London) and Dr Sophie D. Adams (Draviam group) for comments on the manuscript. We also thank Draviam group members for useful discussions.

## Author contributions

D.D. and V.M.D. designed the study. Experiments were performed by D.D. SpinX’s source code was written by D.D. Widefield microscopy dataset generation and data analysis was performed by D.D. Confocal microscopy dataset was generated by C.E. Figures were originally assembled by D.D and edited by C.E, D.D and V.M.D. Annotations were performed by C.E. and D.S. Model benchmarks and testing were performed by C.E., D.S. and D.D. The manuscript was drafted by D.D and edited by C.E. and V.M.D., with comments from N.S. and Draviam group members.

## Ethics declarations

The authors declare no competing financial interests.

