## Supplemental Files for "SpinX: Time-resolved 3D Analysis of Mitotic Spindle Dynamics using Deep Learning Techniques and Mathematical Modelling"

### Supplementary

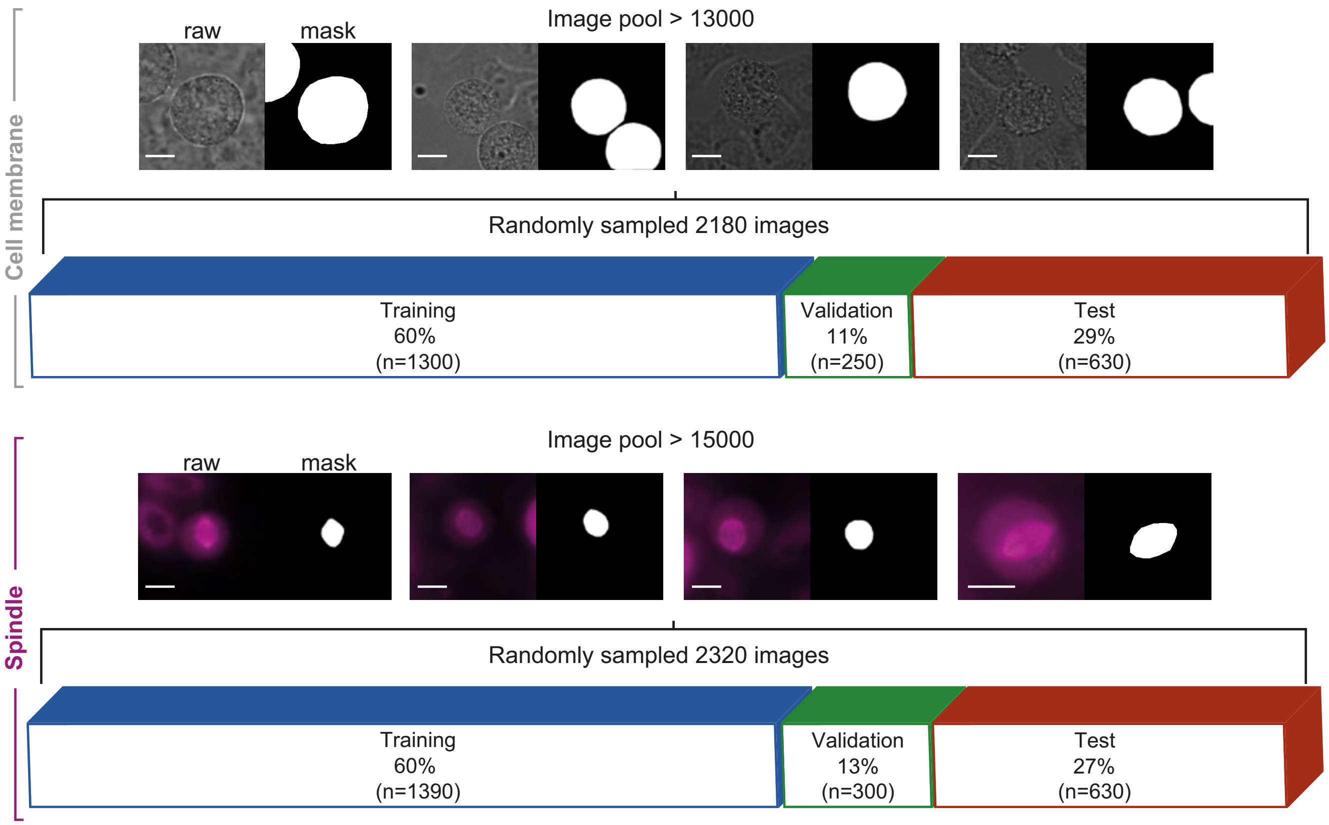


**Supplementary Figure 1. Dataset composition.** Dataset composition of cell membrane (top row) and spindle (bottom row) images with their corresponding masks for training, validation and testing. n corresponds to the number of images that were randomly selected from the image pools. Scale bars: 10 $\mu$m.

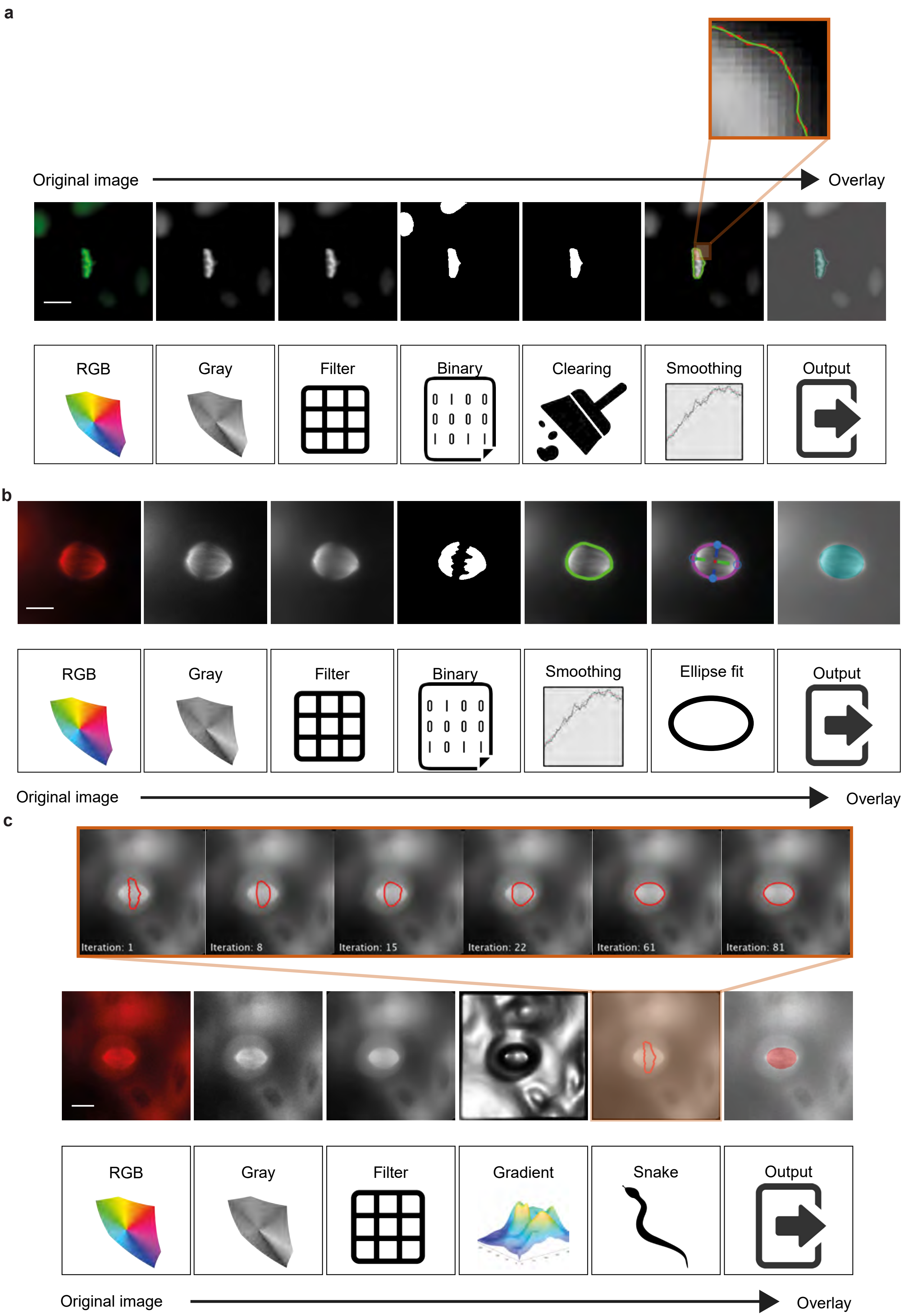
a
Original image
Overlay
RGB
Gray
Filter
Binary
Clearing
Smoothing
Output
b
RGB
Gray
Filter
Binary
Smoothing
Ellipse fit
Output
Original image
Overlay
c
Iteration: 1
Iteration: 8
Iteration: 15
Iteration: 22
Iteration: 61
Iteration: 81
RGB
Gray
Filter
Gradient
Snake
Output
Original image
Overlay

**Supplementary Figure 2.** **SpinX pipelines for automated label generation a.** Conventional image processing pipeline to segment chromosomes. Pipeline includes using a median filter to reduce surrounding noise while preserving information of edges; performing Otsu's thresholding to create a binary image; removing incomplete objects located at the image boundary; extracting boundary pixel information of the metaphase plate and applying a Savitzky-Golay filter to smoothen the boundary. **b.** Conventional image processing pipeline to segment SiR-Tubulin-labelled spindle images. Pipeline includes using a median filter to reduce surrounding noise while preserving information of edges; performing Otsu's thresholding to create a binary image; calculating the binary convex hull image; extracting boundary pixel information of the spindle; applying a Savitzky-Golay signal processing filter for smoothing; and utilising an ellipse fit to obtain the final boundary information. **c.** Conventional image processing pipeline to segment the mCherry-Tubulin-labelled spindle images. Pipeline includes applying a Gaussian filter to reduce surrounding noise while preserving information of edges; calculating the image gradient; using the segmentation mask of the metaphase plate to initiate the inverse snake; extracting boundary pixel information of the spindle and utilising an ellipse fit to obtain the final boundary information. Scale bars: 10 $\mu$m.

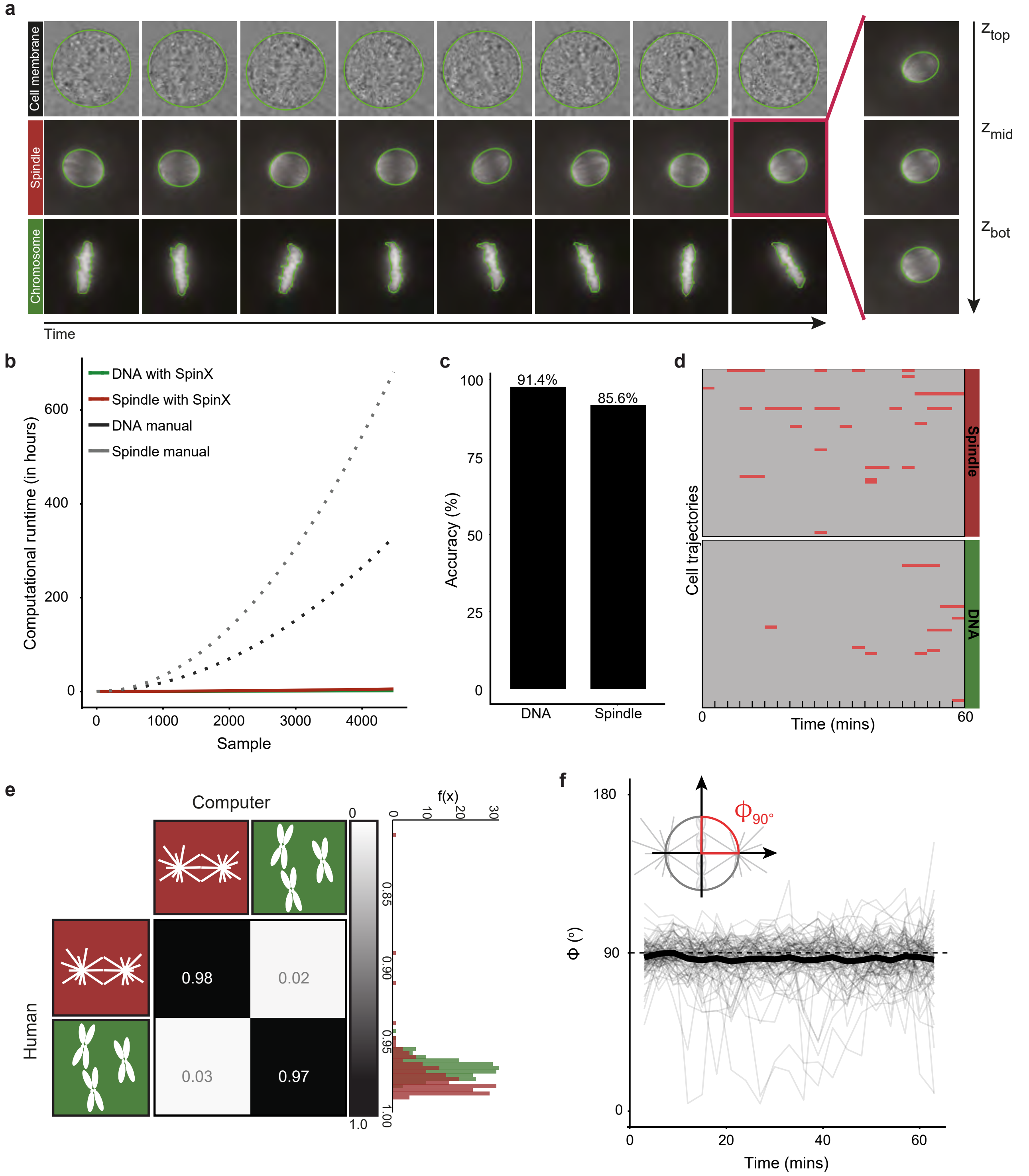

a
Cell membrane
Spindle
Chromosome
Time
Z top
Z mid
Z bot
b
DNA with SpinX
Spindle with SpinX
DNA manual
Spindle manual
Computational runtime (in hours)
Sample
c
Accuracy (%)
91.4%
85.6%
DNA
Spindle
d
Cell trajectories
Spindle
DNA
Time (mins)
e
Computer
Human
0.98
0.02
0.03
0.97
f(x)
f
Φ (°)
Φ90°
Time (mins)

**Supplementary Figure 3. Evaluation of automated label generation with SpinX. a.** Time-lapse image stills of cell membrane, spindle (SiR-Tubulin) and chromosomes. The segmentation results are outlined in green. **b.** Graph shows computational run time for chromosome (green) and spindle (red) versus manual segmentation (black and gray dotted-line), respectively. An exponential function of second order was fitted to predict the remaining time points. **c.** Bar graph shows image-wise accuracy for chromosome and spindle channels. An image is defined as missegmented if SpinX fails to segment the full object's boundaries or if the properties (e.g. orientation) are incorrect. Accuracy percentages are calculated by the number of missegmented frames over the total number of frames. **d.** Automated chromosome and spindle segmentation of cell trajectories over time without correction. Missegmented frames are highlighted in red. The time interval between each frame is 3 minutes. **e.** The correlation matrix displays the calculated correlation coefficient $\rho$ which denotes the matching between human segmentation versus automated segmentation. The histogram shows the distribution of $\rho$ grouped by channels. **f.** Time series graph shows chromosome-spindle formation at metaphase. Dash-line at angle $\phi = 90^{\circ}$ represents perfect perpendicular properties between spindle and chromosomes. N = 4473 images per channel. Data obtained from 88 movies of metaphase cells over at least 6 independent experiments.

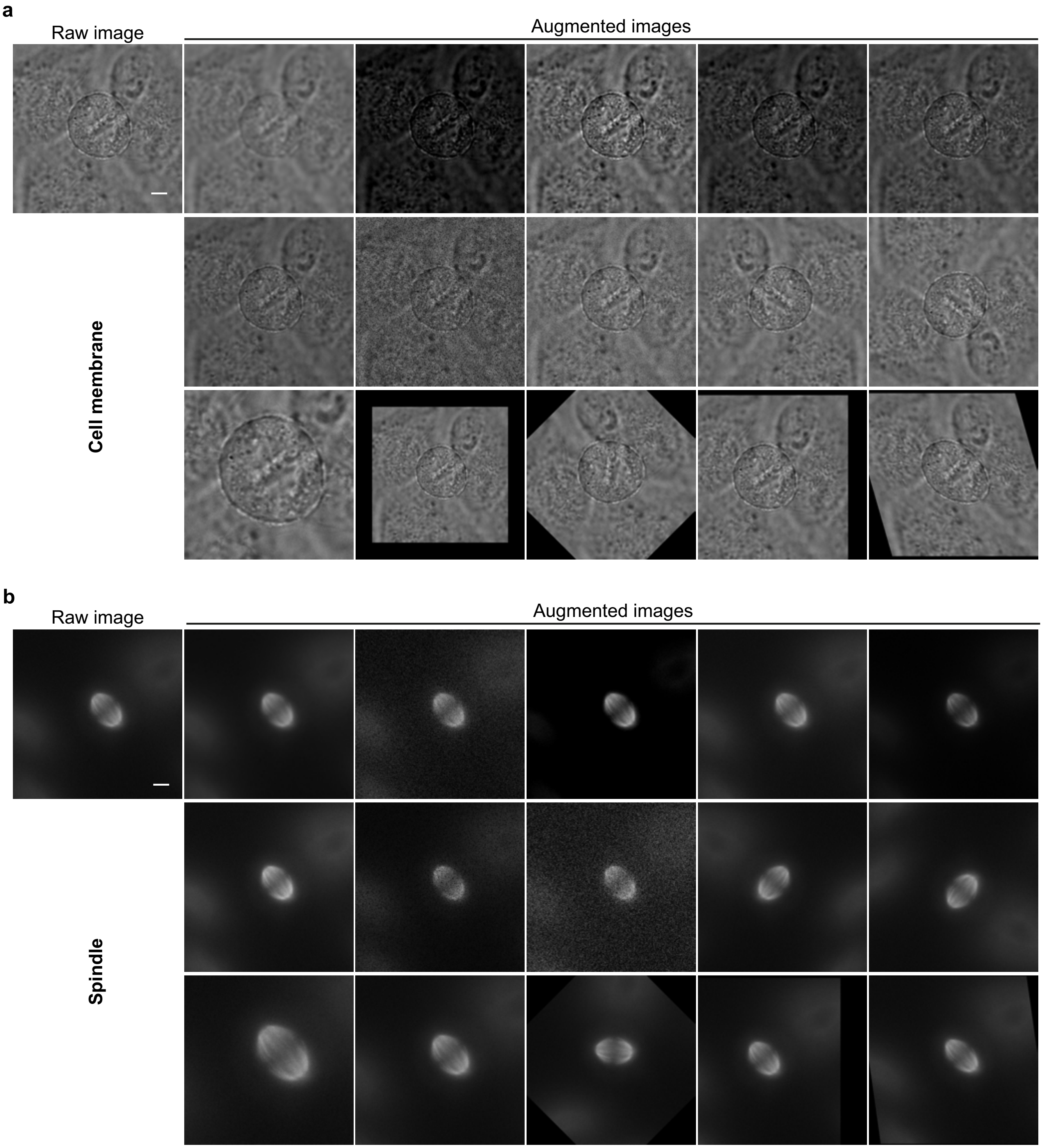


**Supplementary Figure 4. Data augmentation. a.** The data augmentation techniques applied for the cell cortex dataset include (left to right): Gaussian blur, element-wise addition, contrast normalisation, simple pixel value addition, gamma adjustment, multiplying pixel values, dropout pixels, adding Gaussian noise, flipping left/right, flipping up/down, cropping, scaling, rotating, translating, and shearing the image. **b.** The same augmentation techniques as described in (a) were applied for the spindle dataset. Scale bar: 5 $\mu$m.

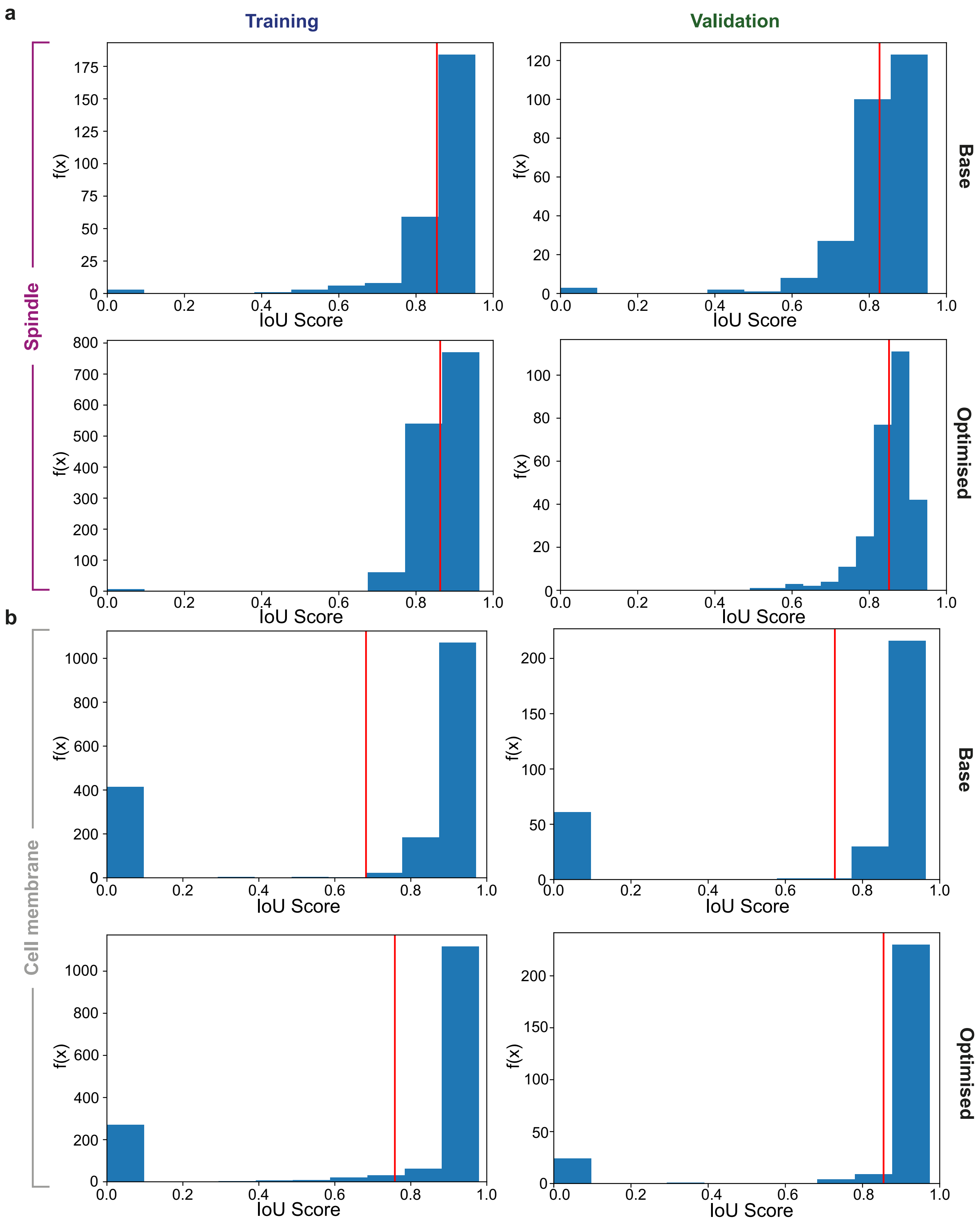


**Supplementary Figure 5. Computational evaluation of SpinX-base and SpinX-optimised. a.** Histogram of IoU for base and optimised models derived from the spindle training and validation datasets. **b.** Histogram of IoU for base and optimised models derived from cell membrane training and validation datasets.

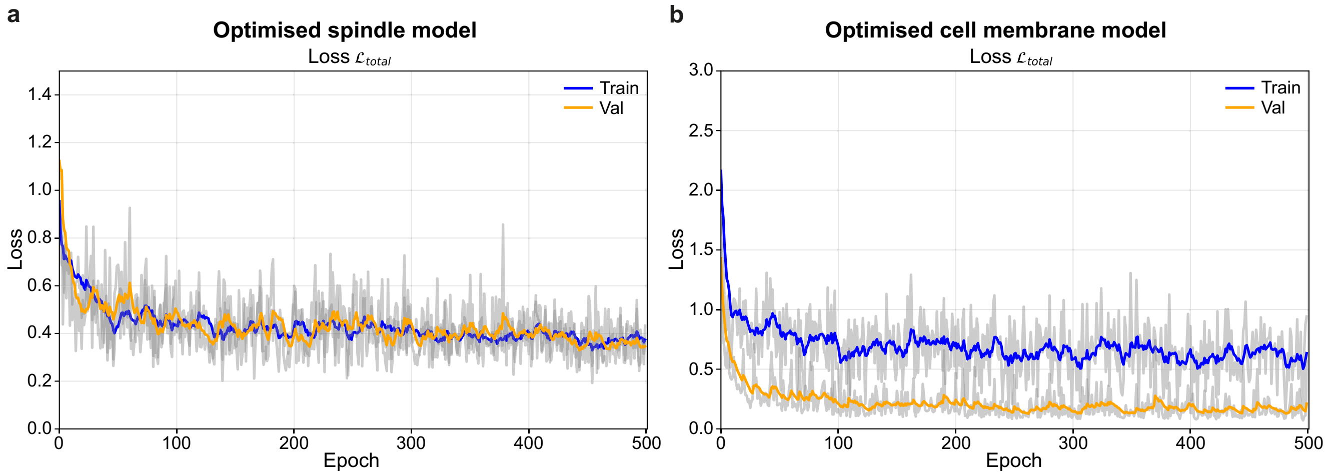


**Supplementary Figure 6. Computational evaluation of SpinX-base and SpinX-optimised. a.** Line graphs show declining training (blue) and validation (orange) Loss functions for the optimised spindle model. **b.** Line graphs show declining training (blue) and validation (orange) Loss functions for the optimised cell membrane model.

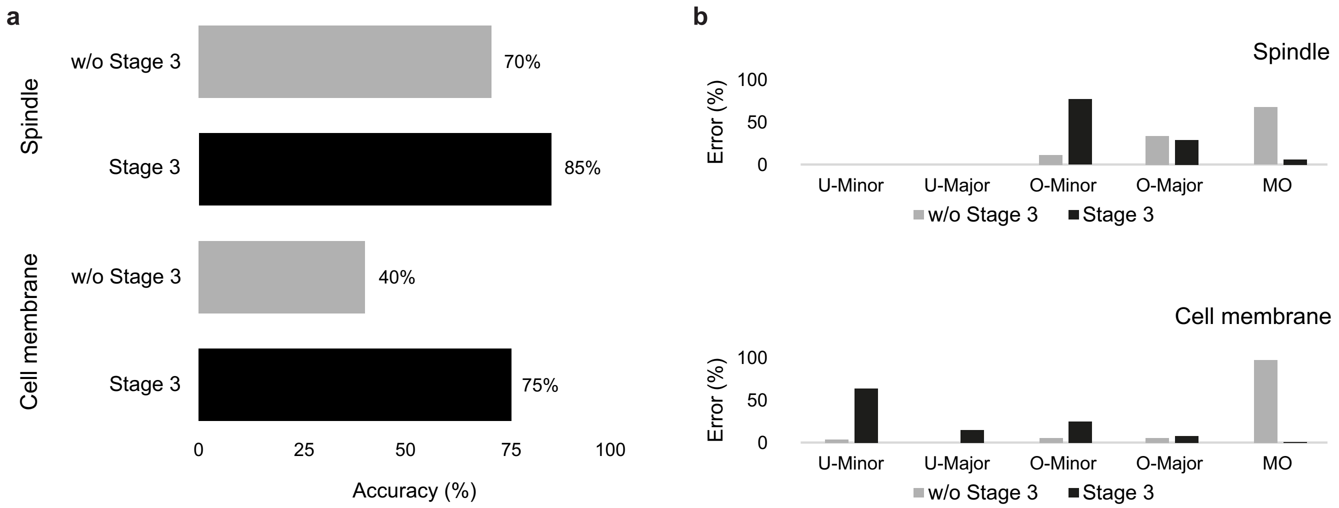


**Supplementary Figure 7. SpinX stage 3 evaluation. a.** Bar chart shows accuracy without (gray) and with stage 3 (black) of SpinX's architecture for spindle and cell membrane models. **b.** Wrongly predicted images from spindle and cell membrane models were further analysed using our error classification system (described in Fig. 2c). n=100 randomly selected images from our image pool were used for studies without stage 3. n=630 images from 10 randomly selected 3D time-lapse movies were used for studies with stage 3.

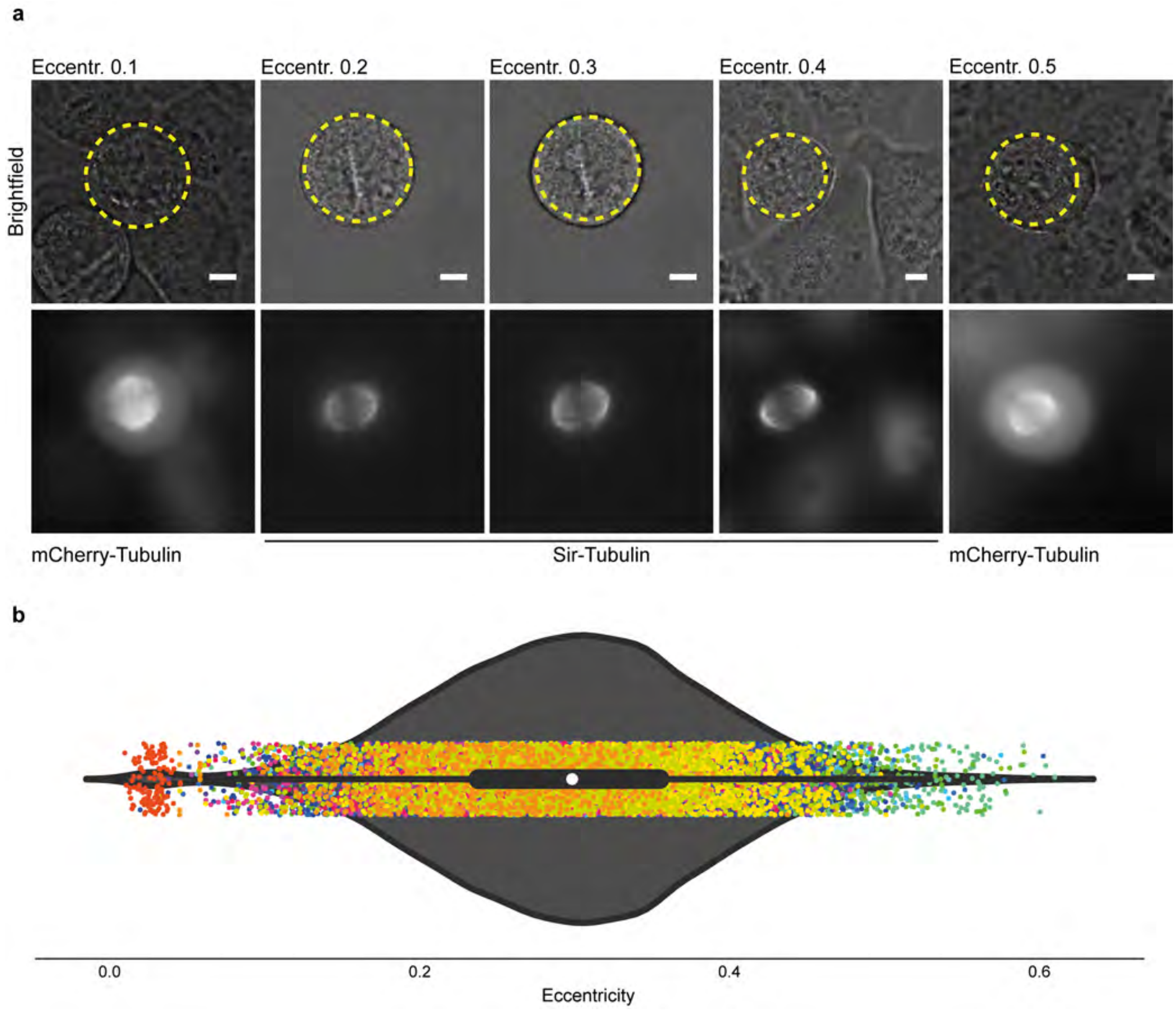


**Supplementary Figure 8. Cell cortex is not fully circular. a.** Representative brightfield images (first row) of HeLa cell lines expressing mCherry-Tubulin or Sir-Tubulin. Cells were selected based on measurements of eccentricity. Intact spindles of the individual cells are represented in the second row. Circular shape with eccentricity = 0 is highlighted in yellow. **b.** Violin plot shows distribution (dots) of eccentricity across 96 3D live-cell movies from 16 experiments. White marker within the box refers to the median, the shaded area refers to the estimated kernel probability density and the box indicates the interquartile range of the data. Measurements from the same dataset share the same color. Scale bar: 5 $\mu$m.

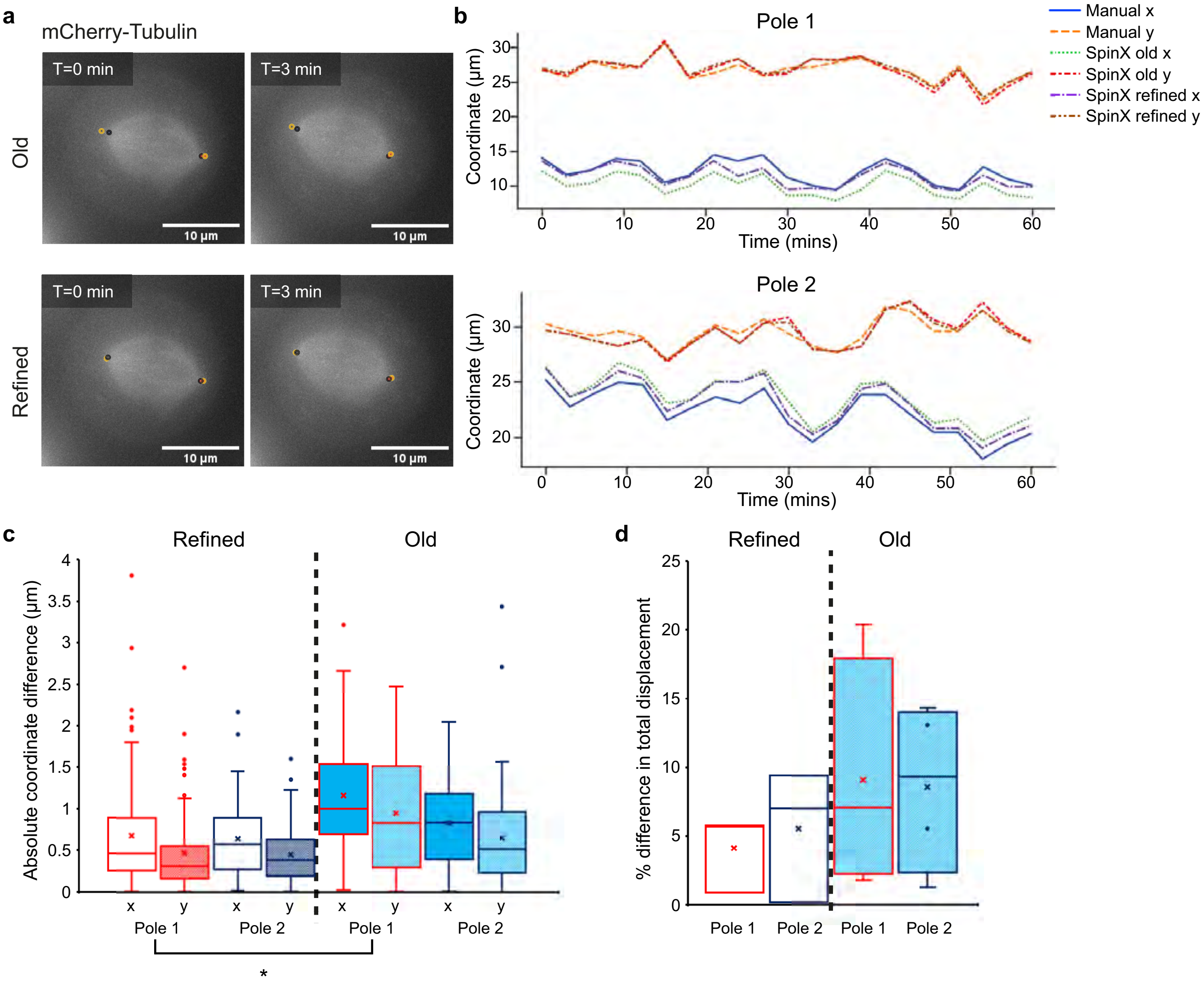


**Supplementary Figure 9. SpinX's refined algorithm for recording pole positions is more reliable compared to its previous iteration. a.** Representative maximum projection images of the mitotic spindle in a HeLa mCherry-Tubulin cell showing pole positions as recorded from SpinX's old (top) and refined (bottom) algorithm. Black circles correspond to the poles recorded by manual analysis, while orange circles correspond to the poles recorded by SpinX. Pole alignment was kept consistent throughout measurements i.e. red-filled circles correspond to pole 1, whereas blue-filled circles correspond to pole 2. T corresponds to the time at which the image was taken. Scale bar = 10 $\mu$m. **b.** Representative traces of the x-y coordinates of the poles of a single cell across time as recorded from manual analysis, and SpinX's old and refined algorithm (Pole 1 - top; Pole 2 - bottom.) **c.** Box plots quantifying the absolute coordinate (x-y) difference between manual analysis and either refined (left) or old (right) SpinX for both Poles 1 and 2. N = 4 cells, each cell consisting of 21 measurements for each pole. Kruskal Wallis H test + post-hoc Dunn's test with Bonferroni adjustment: * $p < 0.05$ Pole 1 refined vs Pole 1 old. **d.** Box plots quantifying the % difference in spindle pole total displacement between manual analysis and either the refined (left) (N=3 cells) or old (right) (N=4 cells) versions of SpinX. The comparison between manual analysis and SpinX was performed based on x-y coordinates alone, but it should be noted that SpinX performs 3D reconstruction, thus predicting x-y positions while taking into consideration the *z*-position.

a

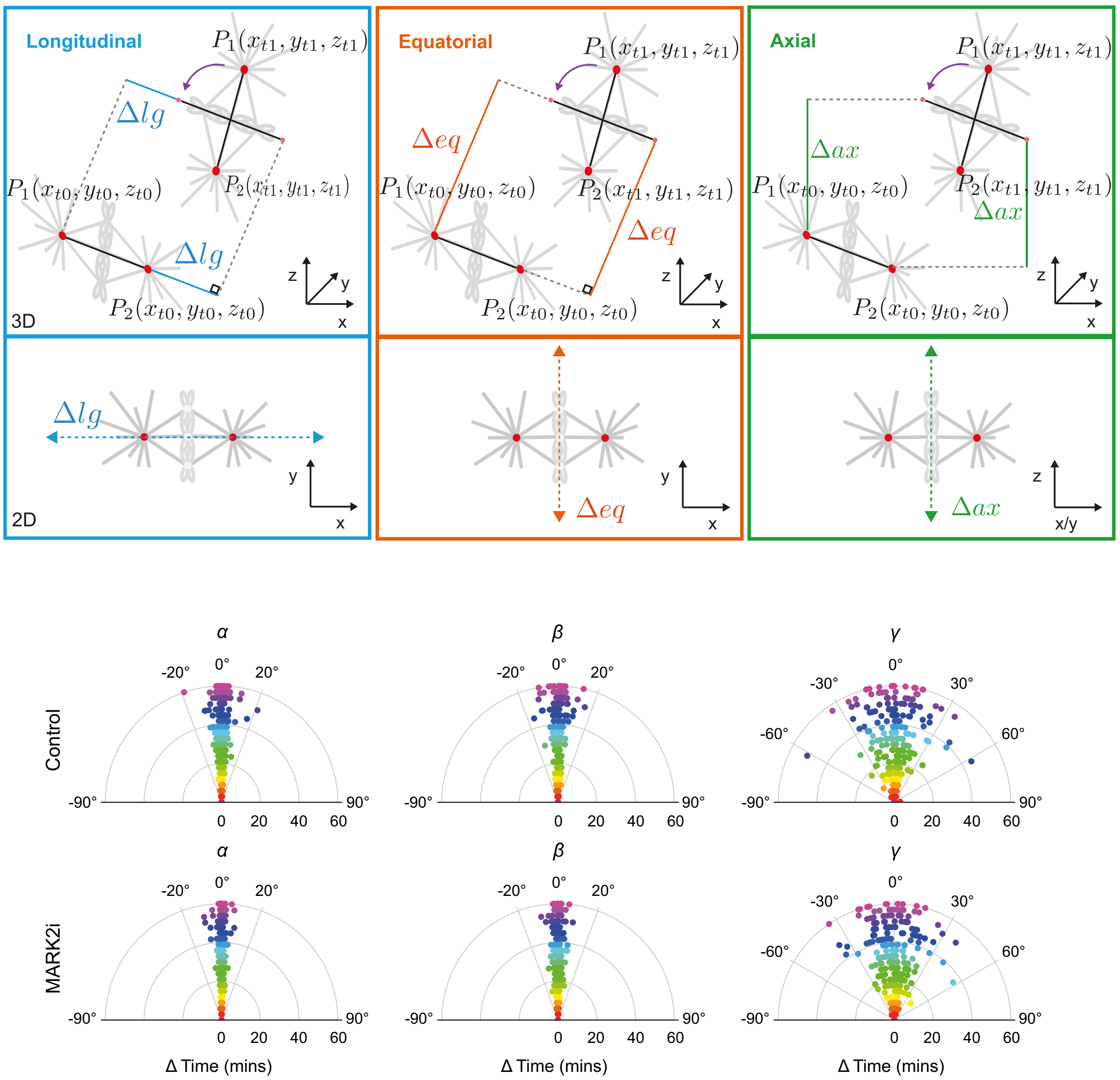


**Supplementary Figure 10. MARK2i does not affect spindle rotation. a.** Illustration shows decomposition of spindle movement (gray) from $t_0$ to $t_1$ in 3D. Red dots correspond to spindle poles $P1$ and $P2$. The spindle length axis is represented by the black line. Purple arc arrow indicates rotational movement of the spindle length axis at $t_1$ when parallel to $t_0$. Blue box: Blue lines cover the longitudinal movement Δlg along the spindle length axis. Orange box: Orange lines cover the equatorial movement Δeq along the spindle width axis which is perpendicular to the spindle length axis. Green box: Green lines cover the axial movement Δax along the spindle axial axis. The 2D representation (bottom row) illustrates the raw decomposed movement. **b.** Polar scatter plot shows spindle rate of rotational movement (in degrees) through time. Measurements at each time point are highlighted with a unique color. $\alpha$, $\beta$ and $\gamma$ angles refer to spindle tumbling, rolling and rotation respectively. Statistical significance was determined by Mann-Whitney U test after a pre-analysis of the underlying distribution with a Shapiro-Wilk test. N=11 control and N=12 MARK2i cells across three experiments.

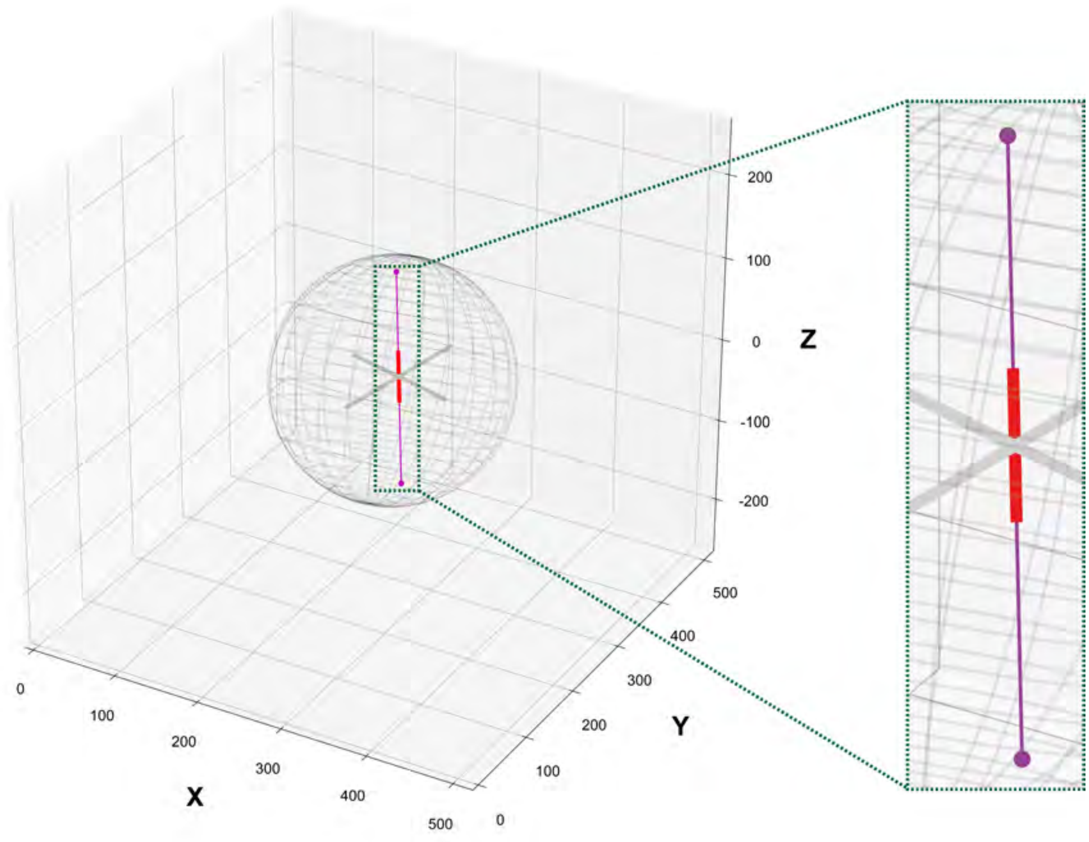


**Supplementary Figure 11. Analytical solution for 3D Ray-tracing.** Mathematically applying the analytical solution (as described in Methods) results to a connecting line (magenta) of two intersection points (i.e. points of contact between the spindle principal axis' endpoints and the cell surface) that overlaps with the spindle height axis (red), hence allowing the calculation of pole-cortex distances in 3D.

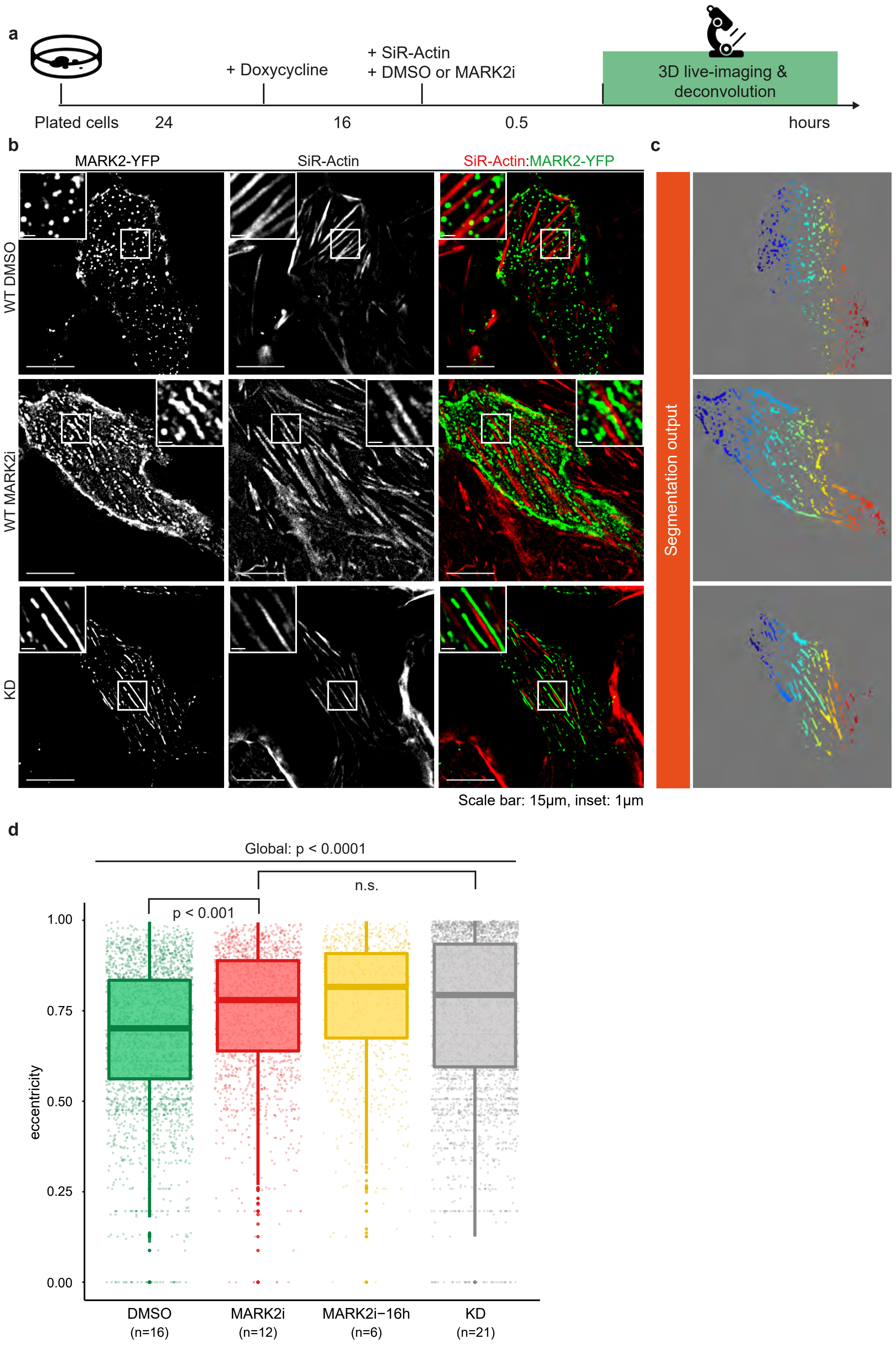
a
+ Doxycycline
+ SiR-Actin
+ DMSO or MARK2i
3D live-imaging & deconvolution
Plated cells
24
16
0.5
hours
b
MARK2-YFP
SiR-Actin
SiR-Actin:MARK2-YFP
WT DMSO
WT MARK2i
KD
c
Segmentation output
Scale bar: 15μm, inset: 1μm
d
Global: p < 0.0001
n.s.
p < 0.001
eccentricity
1.00
0.75
0.50
0.25
0.00
DMSO
(n=16)
MARK2i
(n=12)
MARK2i−16h
(n=6)
KD
(n=21)

**Supplementary Figure 12. MARK2-YFP localises as punctuate-striation pattern upon MARK2 inhibition. a.** Experimental regime and procedures. HeLa cells were exposed to Doxycycline 16 hours before imaging. SiR-Actin (100 nM) and MARK2i (10 $\mu$M) or DMSO (solvent control) were added 30 mins prior to imaging. **b.** Deconvolved $z$-slice of 3D image stacks show MARK2-YFP (green) at the cell-substrate interface in WT treated with DMSO, MARK2i or in KD stained with SiR-Actin dye (red). White boxes show punctate foci pattern in WT (DMSO), punctate-striation pattern in WT (MARK2i) and long striation in KD. **c.** Segmentation results of MARK2-YFP signal for quantification after manually outlining interphase cell boundaries. **d.** The boxplots show eccentricity measurements for DMSO, MARK2i, MARK2i prolonged treatment (16 hours) and KD. 0 represents a perfect circle and 1 a perfect line. For global group comparison, a generalised linear model (GLM) was fitted (non-normality was determined with Shapiro-Wilk test) followed by a post-hoc analysis for pair-wise comparison (Dunn's post-hoc test) after Multiple-Comparison Kruskal-Wallis (MCKW) at a significant level of $p < 0.01$. n refers to the number of cells obtained from three experiments. Scale bars: 15 $\mu$m, 1 $\mu$m for inset.

| | Base model | Optimised model |
|---|---|---|
| Annotation | Less experienced | Refined annotations by experts |
| Sample size | 800 (cell mem.)<br>900 (spindle) | 1300 (cell mem.)<br>1390 (spindle) |
| No. Epoch | 200 | 500 |
| Weight decay | 0.0001 | 0.001 |

**Supplementary Table 1. Differences between SpinX-base and SpinX-optimised.** Table shows differences in annotation quality, sample and hyperparameter values between the base and optimised models.

| | | Training | | | Validation | | |
|---|---|---|---|---|---|---|---|
| Type | Model | Loss | AP | mIoU | Loss | AP | mIoU |
| Spindle | Base | 0.2293 | 0.976 | 0.874 | 0.2406 | 0.867 | 0.854 |
| | Optimised | 0.1928 | 0.988 | 0.861 | 0.2634 | 0.920 | 0.852 |
| Cell membrane | Base | 0.1467 | 0.975 | 0.682 | 0.0838 | 0.866 | 0.728 |
| | Optimised | 0.1761 | 0.989 | 0.758 | 0.0712 | 0.990 | 0.850 |

**Supplementary Table 2. Evaluation of SpinX-base and SpinX-optimised models.** Table shows evaluation results of base and optimised models through comparisons between Loss, mean AP and mean IoU computed from training and validation datasets.

| | Error in training annotation | Error validation annotation |
|---|---|---|
| Spindle | 31% (N=900) | 46% (N=300) |
| Cell membrane | 44% (N=800) | 38% (N=250) |

**Supplementary Table 3. Evaluation of annotation.** Table shows percentage of annotations with errors. These images were identified, re-annotated by an expert, and subsequently used for training and validation of the optimised model.

| | | Corrected | | |
|---|---|---|---|---|
| N | Total time frame | Spindle height | Spindle width | Spindle length |
| 10 | 210 | 29%<br>60 out of 210 | 52%<br>109 out of 210 | 48%<br>101 out of 210 |

**Supplementary Table 4. Spindle tracking evaluation.** Spindle tracking evaluation of 10 random live-cell movies. The proportion of corrections were analysed for spindle height, width and length axes.

| **Parameter** | **Description** | **Value** | **Reference** |
|---|---|---|---|
| NA | NA of the objective lens | 1.4/1.42 | DeltaVision |
| $n_s$ | RI of the sample | 1.34 (Cytosol) | [64] |
| $n_i$ | RI of the immersion | 1.522 | DeltaVision |
| $\lambda$ | Wavelength | FITC: 490nm/525nm | DeltaVision |
| | | TRITC: 555nm/605nm | DeltaVision |
| | | mCherry: 572nm/632nm | DeltaVision |
| | | CY5: 645nm/705nm | DeltaVision |
| M | Magnification | 100/160 | DeltaVision |
| $t$ | Working distance | 150 nm | DeltaVision |
| $x_d, y_d$ | Lateral resolution | 0.100 $\mu$m | DeltaVision |
| $z_d$ | Axial resolution | 0.250 $\mu$m | DeltaVision |
| X | PSF array width | 256 | - |
| Y | PSF array height | 256 | - |
| Z | PSF array depth | 128 | - |

**Supplementary Table 5. Parameters used for PSF simulation.**

**Supplementary Video 1. Spindle and cell cortex tracking in 3D with SpinX.** Video shows composite time-lapse movies of spindle movements in a metaphase HeLa cell expressing mCherry-Tubulin (spindle marker in purple). Raw time-lapse image of mCherry-Tubulin labelled spindle (purple) merged with corresponding brightfield (grey) image of the cell (top-left) or SpinX's AI module predicted cell cortex (outlined in blue (top-middle). Top-right, movie of SpinX's mathematical object modelling output showcasing spindle pole movements in 3D through time within the metaphase cortex, with an inset displaying the 3D reconstructed spindle; bottom, animated graph highlighting the dynamic change in pole-cortex distances of the two spindle poles as tracks through time.
