## Supplementary figures and images for "SpinX: Time-resolved 3D Analysis of Mitotic Spindle Dynamics using Deep Learning Techniques and Mathematical Modelling"

### Supplementary Movie

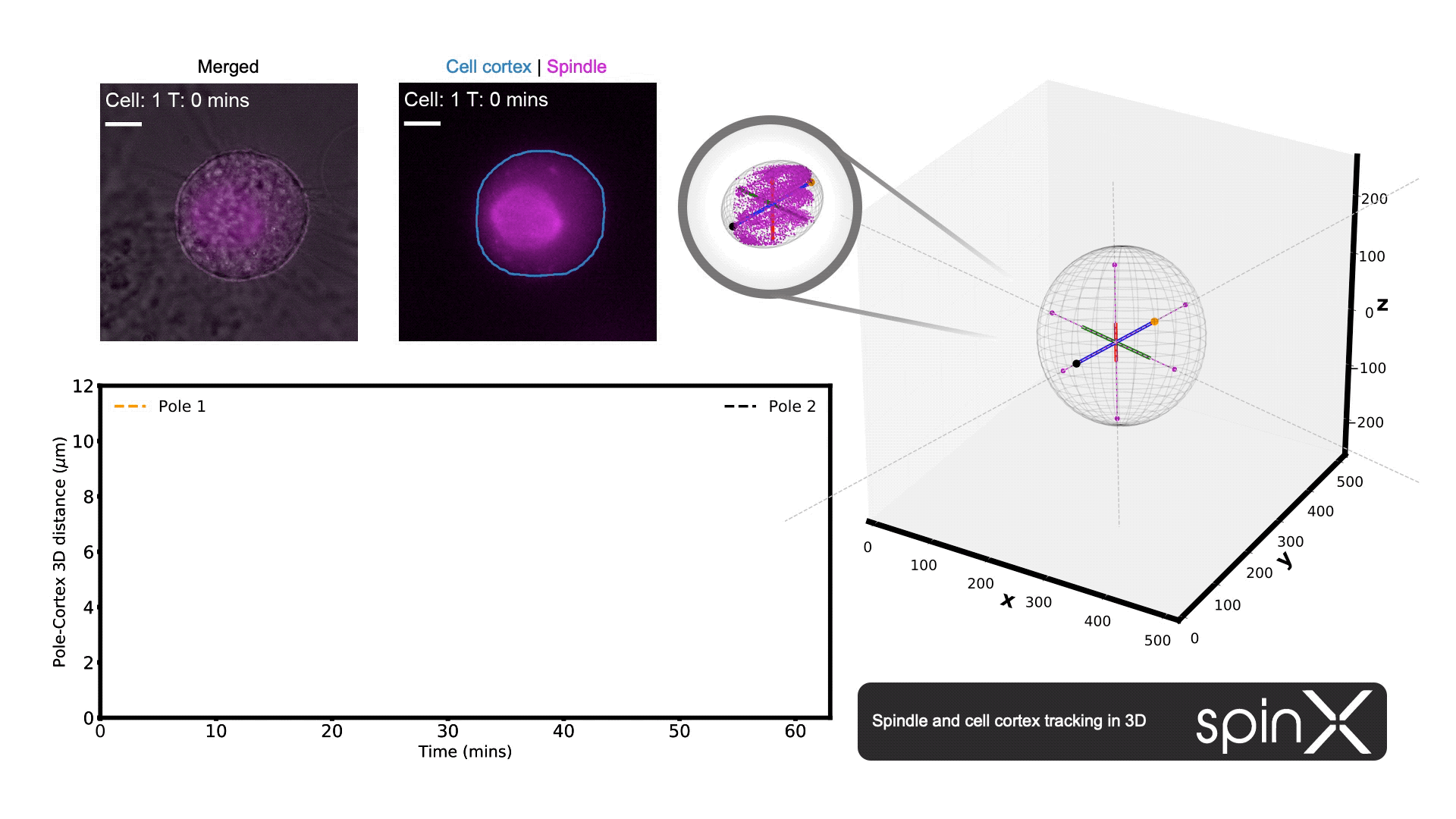
